# Comparison of decision-related signals in sensory and motor preparatory responses of neurons in Area LIP

**DOI:** 10.1101/169219

**Authors:** S. Shushruth, Mark Mazurek, Michael N. Shadlen

## Abstract

Neurons in the lateral intraparietal area (LIP) of Macaques exhibit both sensory and oculomotor preparatory responses. During perceptual decision making, the preparatory responses have been shown to track the state of the evolving evidence leading to the decision. The sensory responses are known to reflect categorical properties of visual stimuli, but it is not known if these responses also track evolving evidence. We compared sensory and oculomotor-preparatory responses in the same neurons during a direction discrimination task when either the discriminandum (random dot motion) or an eye movement choice-target was in the neuron’s response field. Both configurations elicited task related activity, but only the motor preparatory responses reflected evidence accumulation. The results are consistent with the proposal that evolving decision processes are supported by persistent neural activity in the service of actions or intentions, as opposed to high order representations of stimulus properties.

**SIGNIFICANCE STATEMENT:** Perceptual decision making is the process of choosing an appropriate motor action based on perceived sensory information. Association areas of the cortex play an important role in this sensory-motor transformation. The neurons in these areas show both sensory- and motor-related activity. We show here that, in the macaque parietal association area LIP, signatures of the process of evidence accumulation that underlies the decisions are predominantly reflected in the motor-related activity. This finding supports the proposal that perceptual decision making is implemented in the brain as a process of choosing between available motor actions rather than as a process of representing the properties of the sensory stimulus.

## INTRODUCTION

The life of animals is a constant process of deciding what to do next based on, among other things, the perception of the world around them. In primates, perceptual decision making has evolved into an efficient mechanism of translating the perceived state of the world into possible motor actions (Cisek & Kalaska 2005, Klaes et al 2011, Kubanek & Snyder 2015). The motor system receives continuous access to evolving perceptual decisions and maintains a graded level of preparedness based on the quality of the incoming evidence (Gold & Shadlen 2000, Selen et al 2012). This sensorimotor transformation is particularly evident in the parietal and prefrontal association cortices, where neurons encoding the motor actions associated with the choices on offer also represent evolving decisions (Bollimunta & Ditterich 2011, de Lafuente et al 2015, Ding & Gold 2012, Kim & Shadlen 1999, Roitman & Shadlen 2002). Thus, perceptual decision making can be framed as a choice between available motor actions (Cisek 2007, Cisek & Kalaska 2010, Shadlen et al 2008).

Yet, perceptual decisions do not feel like they are about potential actions but about propositions or stimulus properties. Indeed, one can make a decision without knowledge of the action that will be required to act on it. In such situations, one might expect neural circuits involved in motor planning to be irrelevant to the decision process (Gold & Shadlen 2003). However, it has been shown that even then, neurons in the parietal association areas carry a representation of the properties of the stimulus that will be relevant for future actions (Bennur & Gold 2011, Freedman & Assad 2006, Goodwin et al 2012). It is possible that such an ‘abstract’ representations of decision relevant information—independent of the possible motor actions—coexist with representations of decisions as intended actions (Freedman & Assad 2011). Whether such simultaneous representations exist in the same association area has not been investigated before. Consequently, it is also not known if such abstract representations play a role in the decision-making process.

We used the random-dot motion (RDM) direction discrimination task (Newsome et al 1989) to investigate these questions. In this task, the animals discern the net direction of a stochastic motion stimulus and report their decision by making a saccade to one of two choice targets that is along the direction of the perceived motion. This task is particularly well suited for our purposes. First, optimal performance on this task demands integration of motion evidence over time. This prolonged deliberation time allows characterization of whether a neural population is participating in the process of evidence accumulation or not. Second, there exists a theoretical framework—bounded accumulation of noisy evidence to a decision threshold (aka drift-diffusion, Palmer et al 2005, Smith & Ratcliff 2004)— that accounts quantitatively for the speed and accuracy of decisions in this task. Third, it has been shown that responses of neurons in several areas of the brain involved in planning saccadic eye movements represent the evolving decision in this task (Ding & Gold 2010, Ding & Gold 2012, Horwitz & Newsome 1999, Kim & Shadlen 1999, Shadlen & Newsome 1996).

We focused on the parietal sensorimotor association area LIP. Many neurons in LIP respond to both the presence of a sensory stimulus in, and to a planned saccade into their response fields (Barash et al 1991b). We recorded the responses of the same set of neurons during the RDM discrimination task in two configurations — when the response field contained the RDM stimulus and when it contained one of the choice targets. We show that the neurons represent the moment-by-moment accumulation of sensory evidence only in the latter configuration, that is, when they are involved in the planning of the motor action required to report the choice.

## MATERIALS AND METHODS

All training, surgery, and experimental procedures were conducted in accordance with the National Institutes of Health *Guide for Care and Use of Laboratory Animals* and were approved by the University of Washington Institutional Animal Care and Use Committee (IACUC Protocol # 2896-01).

### Experimental Design and Statistical Analysis

#### Neural recordings

We recorded activity of 49 well isolated single units from area LIPv (Lewis & Van Essen 2000) of two adult female rhesus monkeys (*Macaca mulatta*) trained on the random-dot motion direction discrimination task. MRI was used to localize LIPv and to target recording electrodes. Within this putative LIPv, we screened for neurons that had both visual responses and spatially selective persistent activity. The persistent activity was assessed using a memory-guided saccade task (Gnadt & Andersen 1988). In this task, a target is flashed in the periphery while the monkey fixates on a central spot. The monkey has to remember the location of the target and execute a saccade to that location when instructed. The response field (RF) of each neuron was identified as the region of visual space that elicited the highest activity during the interval between the target flash and the eventual saccade. For the majority of neurons in LIPv, this region also elicits the strongest visual response (Platt & Glimcher 1998). During the recording sessions, visual and persistent activities were assessed qualitatively. We confirmed these properties by analyzing the following responses acquired during the experiment: (*i*) the response to RDM presented in the RF, 100-300 ms after onset and (*ii*) delay period activity, 100-300 ms before a saccade into the RF. We confirmed that both proxies were greater than baseline activity, 0-200 ms before the appearance of a visual stimulus in the RF.

#### Behavioral Task

The choice-reaction time direction discrimination task is similar to previous studies (Roitman & Shadlen 2002). The animal initiates a trial by fixating on a point (fixation point; FP) presented on an otherwise black screen. Two choice-targets then appear on the screen. After a variable delay (drawn from an exponential distribution of mean 750 ms), the random-dot motion (RDM) stimulus is displayed in an imaginary aperture (i.e., invisible borders) of 5°-9° diameter at a third location. The first three frames of the stimulus consist of white dots randomly plotted at a density of 16.7 dots • deg^−2^ • s^−1^. From the fourth frame, each dot from three frames before is replotted — either displaced in one direction along the axis connecting the two targets, or at a random location. The probability with which a dot is displaced in the direction of one of the targets determines the stimulus strength (coherence) and on each trial, this was randomly chosen from the set C = [0, 0.032, 0.064, 0.128, 0.256, 0.512]. The motion strengths and the two directions were randomly interleaved. Importantly, the monkey was allowed to view the stimulus as long as it wanted and indicate the perceived direction of motion with a saccade to the target that lay in that direction to obtain a liquid reward. Rewards were given randomly (p=0.5) for the 0% coherence motion condition.

During recording from each isolated neuron, the choice-targets and the RDM were presented in two configurations (Figure 1). In the ‘Target-in-RF’ configuration, one of the choice-targets overlay the neuronal RF. In the ‘RDM-in-RF’ configuration, the RDM stimulus was presented in the RF. The two configurations were alternated in blocks (median block size 90, IQR 60-120). The order of blocks was randomized across neurons (23 started with Target-in-RF blocks; 26 with RDM-in-RF blocks) and each neuron was recorded with at least one block of trials in each configuration. For 33 of the neurons, the targets and the dot stimuli were placed 120° apart on an imaginary circle (as shown in Figure 1). For the remaining 16 neurons (in one monkey), the targets and the dot stimulus were aligned linearly in both configurations. Since the directions of motion varied across sessions, we adopted the following conventions. In the Target-in-RF configuration, the direction of motion towards the target in the RF for each neuron was considered the ‘positive’ direction. In the RDM-in-RF configuration, the positive direction was assigned *post hoc* from the neural recordings: the direction of motion that elicited the higher mean response.

**FIGURE 1:**
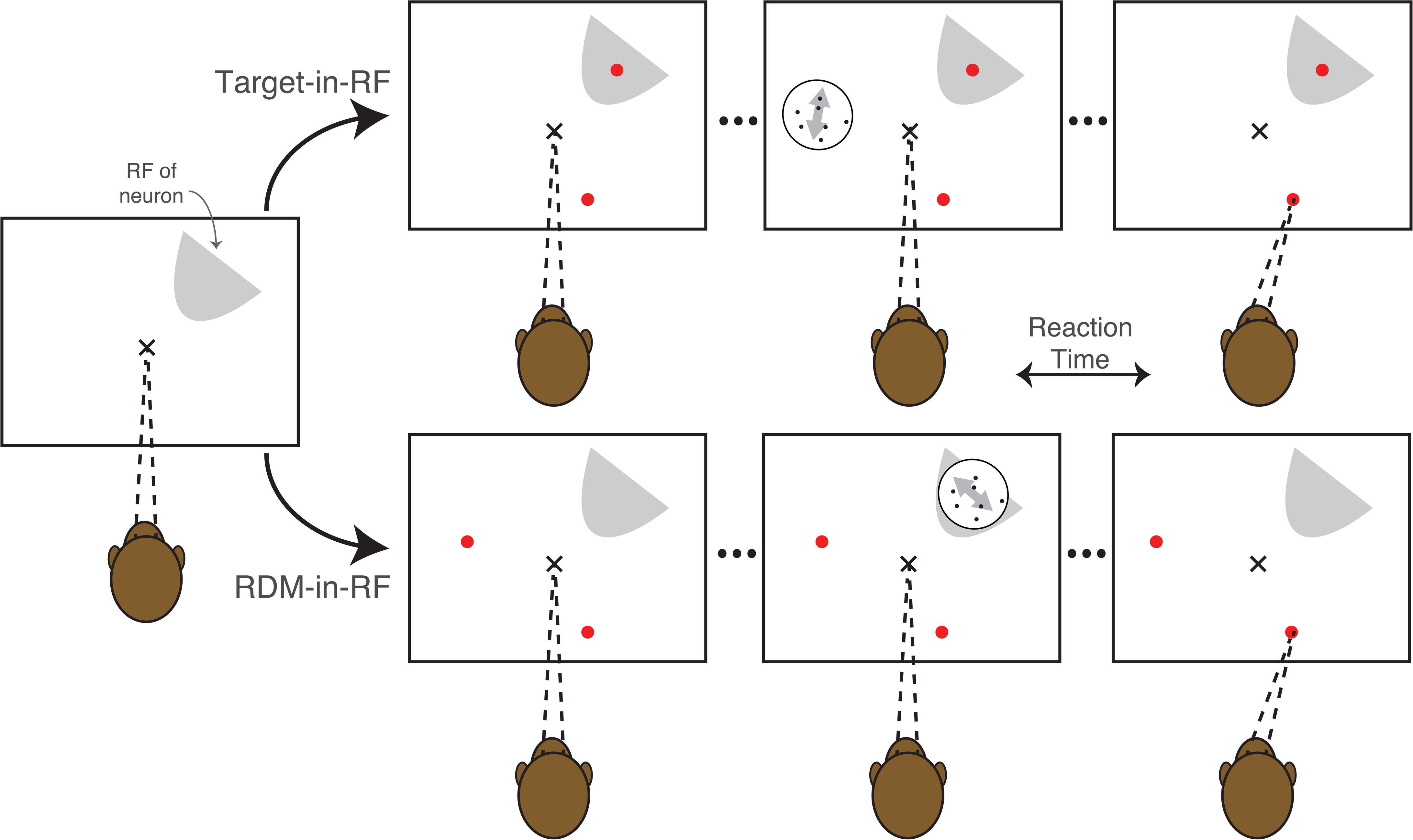
Behavioral task configurations. The monkey fixates at an instructed location (x) and then two choice targets (red dots) appear in one of two configurations:(1) *Target-in-RF*: One of the targets is situated in the RF of the neuron being recorded from, and (2) *RDM-in-RF*: Both targets are situated outside the RF. In the next step, the RDM is presented either inside (RDM-in-RF) or outside the RF (Target-in-RF). The monkey is free to report its decision any time after the appearance of the RDM by making a saccade to one of the targets.

All statistical tests are described in the pertinent sections of Materials and Methods.

### Analyses of behavioral data

The accuracy and reaction times (RT) of the monkeys were fit by a bounded evidence accumulation model (Shadlen et al 2006). In the parsimonious application of this model employed here, the instantaneous evidence about motion at each time step is assumed to arise from a normal distribution with variance *Δt* and mean *κ*(*C* − *C*_%_)Δ*t*, where *C* is the signed motion coherence, *C0* is a bias, and *κ*, a scaling parameter. This instantaneous evidence is accumulated over time and the decision process terminates when the accumulated evidence reaches one of the bounds ±*B* leading to the choice of one of the targets. The mean RT is the expectation of the time taken for the accumulated evidence to reach the bound plus a constant — the non-decision time *tnd* comprising sensory and motor delays. To account for asymmetric reaction times in some configurations, we used two different non-decision times (*tnd1* and *tnd2*) for the two target choices. In this framework, the mean RT for the correct choices (i.e. choices consistent with the sign of the drift rate, *κ*[*C* − *C*_%_]) is described by

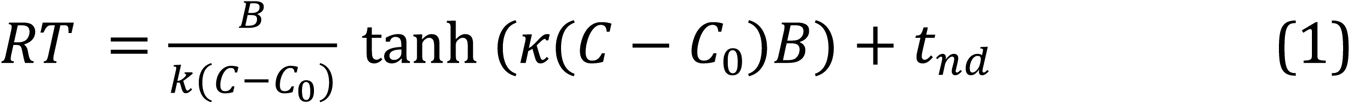

Further, the choice distributions are described by

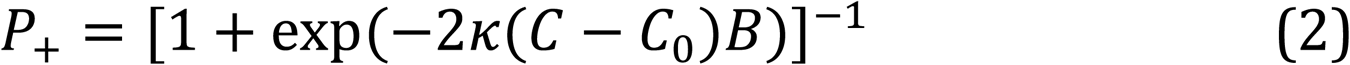

where *P*_+_ is the probability of choosing the target consistent with the ‘positive’ direction of motion. We fit Equation 1 to the RT data and used the fitted parameters to predict the choice functions (Equation 2) (Gold & Shadlen 2002, Kang et al 2017). We first established an estimate of *C0* from a logistic fit of the choices. Because the parsimonious model explains only the RT when the choice is consistent with the sign of the drift rate (Ratcliff & Rouder 1998), we used the mean RT for positive choices at *C–C0>*0 and negative choices *for C–C0*<0. We then fit *κ, B, tnd1* and *tnd2* and used the values of *κ* and *B* in Equation 2 to establish predictions of choice (Figure 2).

**FIGURE 2:**
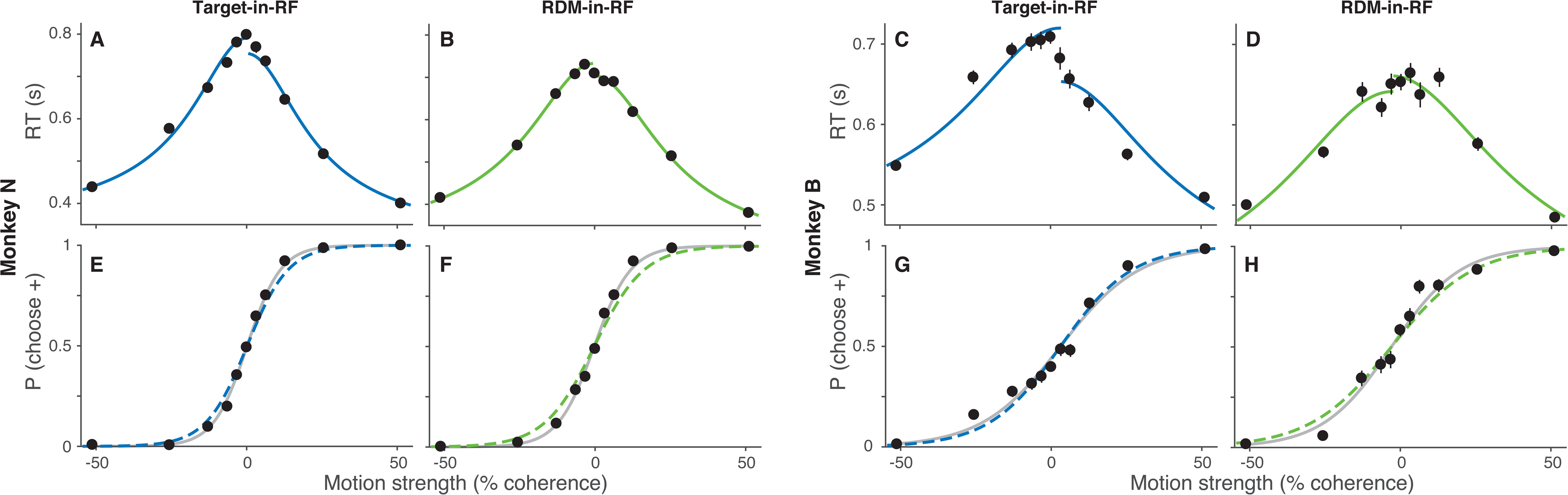
Predicting choices from diffusion-to-bound models fit to RTs. ***A-D*:** RTs of the two monkeys as a function of motion strength in the two task configurations (see Methods for convention on sign of motion strength). Solid lines show the fits of a diffusion-to-bound model. Data includes the trials at 0% motion strength in which the monkey chose the target consistent with its bias (established from logistic fits to the choice data) and correct trials at other motion strengths. ***E-H*:** The probability the monkey chooses the target consistent with positive motion direction, plotted as a function of motion strength. The dashed lines are predictions from the corresponding fits of the RTs. Gray lines are fits to the choice data (logistic regression).

We evaluated the fidelity of these predictions by comparing the predictions to a logistic regression fit of the choice data. To demonstrate that these predictions were not a trivial result of monotonic ordering of RTs by motion strength, we compared them to predictions from 10,000 pseudorandomly generated RT vs. coherence functions that preserved the order of RTs. To generate these functions, we retained the observed RTs for the minimum (−51.2%), maximum (+51.2%) and 0% coherences and used ordered random values within this range for the other coherences. We quantified the magnitude of the perturbation as the average of the percentage change from the observed RT at each coherence. We then performed the steps above to fit these perturbed RTs to establish a new predicted choice function. We estimated the probability of obtaining a predicted choice function as good or better that the ones derived from data as a function of the size of the perturbation. We report the minimal perturbation at which p<0.01.

To obtain a more precise estimate of decision times, we fit an elaborated version of the bounded evidence accumulation model (Extended data Figure 2-1) simultaneously to both choices and reaction times (including both correct and error trials). In this model, the decision bounds (*B*) collapse with time (*t*) such that

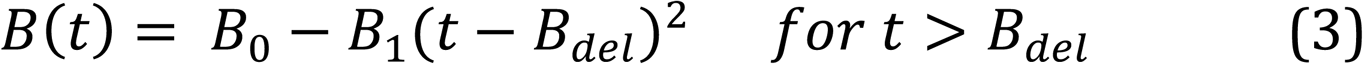

where *B0* is initial bound height, *B1* is the rate of collapse and *Bdel*, the delay to onset of collapse. The non-decision time is modeled as a normal distribution with mean *tnd* and standard deviation *σtnd*. A separate non-decision time was used for decisions terminating at each of the two bounds. This model was fit by maximizing the log likelihood of the observed responses (choice and RT) on each trial to numerical solutions for the probability densities of terminating at *±B*(*t*) (Churchland et al 2008, Kang et al 2017). The mean decision times were obtained from these fits and their standard error estimated from fitting the model to resampled trials (i.e., the standard deviation of the means from 100 iterations).

### Analyses of neural data

Population responses were computed as the average of all trials from all neurons after smoothing each trial with a 75 ms wide boxcar filter (Figure 3A-D). The smoothing was only for visualization and all analyses were conducted on the raw spike data (1 ms resolution). To visualize the coherence dependent buildup of activity (Insets of Figure 3A,C), we detrended individual neuronal responses by subtracting the average responses across all coherences for the same neuron (separately for each task configuration).

**FIGURE 3:**
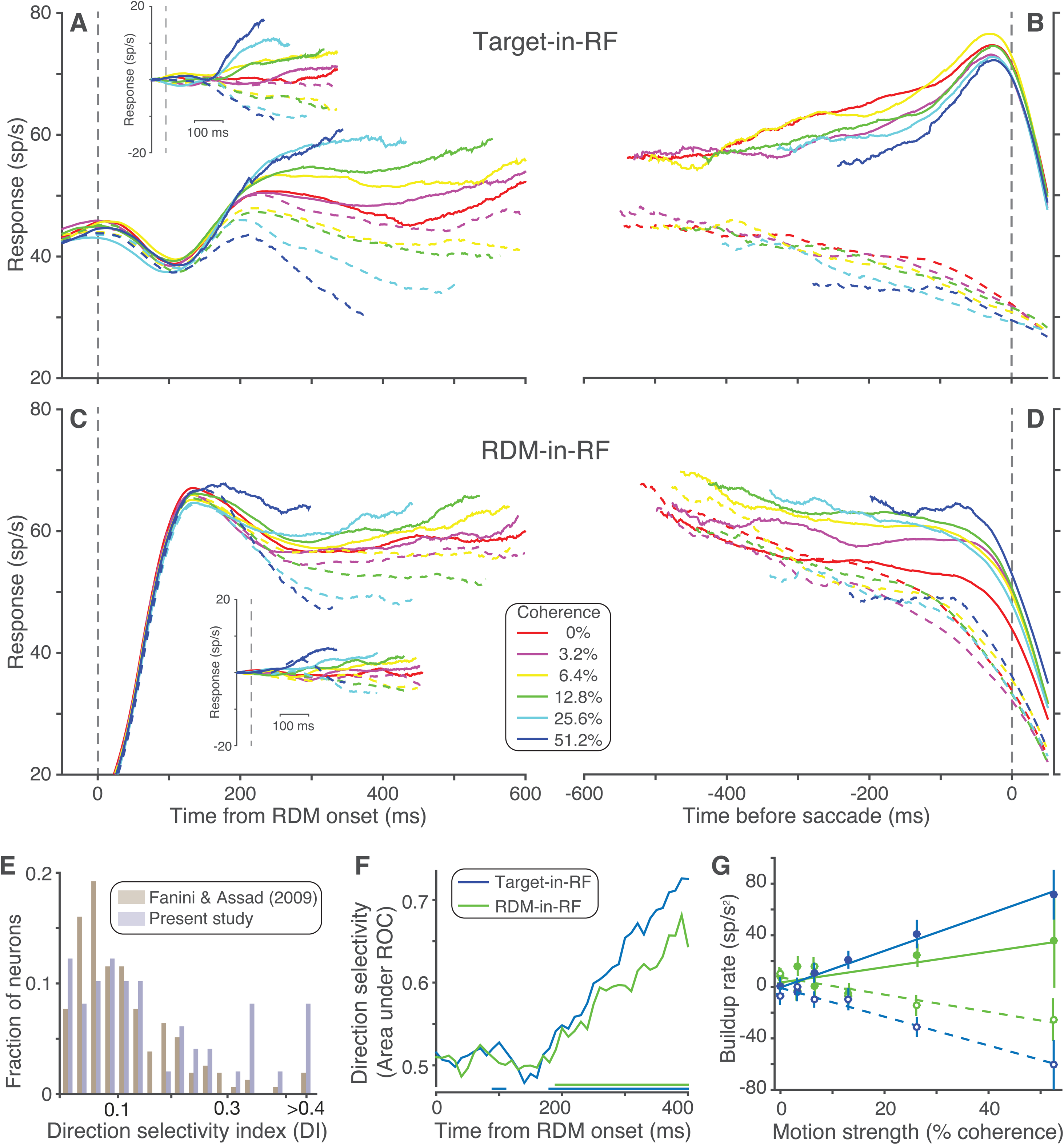
Neural population responses. Average response of the recorded neural population during Target-in-RF ***(A,B)*** and RDM-in-RF ***(C,D)*** configurations. Panels ***A,C*** are aligned to the onset of RDM and include all trials sorted by direction and strength of motion. Insets show average of detrended responses (i.e., after subtraction of the mean response for all motion strengths, for each neuron). Panels ***B,D*** are aligned to the saccade and includes correct trials (and 0% coherence trials sorted by the animal’s choices). ***E:*** Histograms of the distribution of Direction Selectivity Index (DI) for the neural population recorded by Fanini and Assad (2009) and for the neural population in the RDM-in-RF configuration of the present study. ***F:*** Area under ROC for responses to the two directions of motion at 51.2% coherence computed in 40 ms bins. The colored lines at the bottom indicate the time bins in which this metric was significantly >0.5 for the corresponding configuration. ***G:*** The relation between the response buildup rate and motion strength. Filled circles are data from trials with motion in the neuron’s preferred direction and unfilled circles for the opposite motion direction. Solid and dashed lines are corresponding linear regression model fits.

**FIGURE 4:**
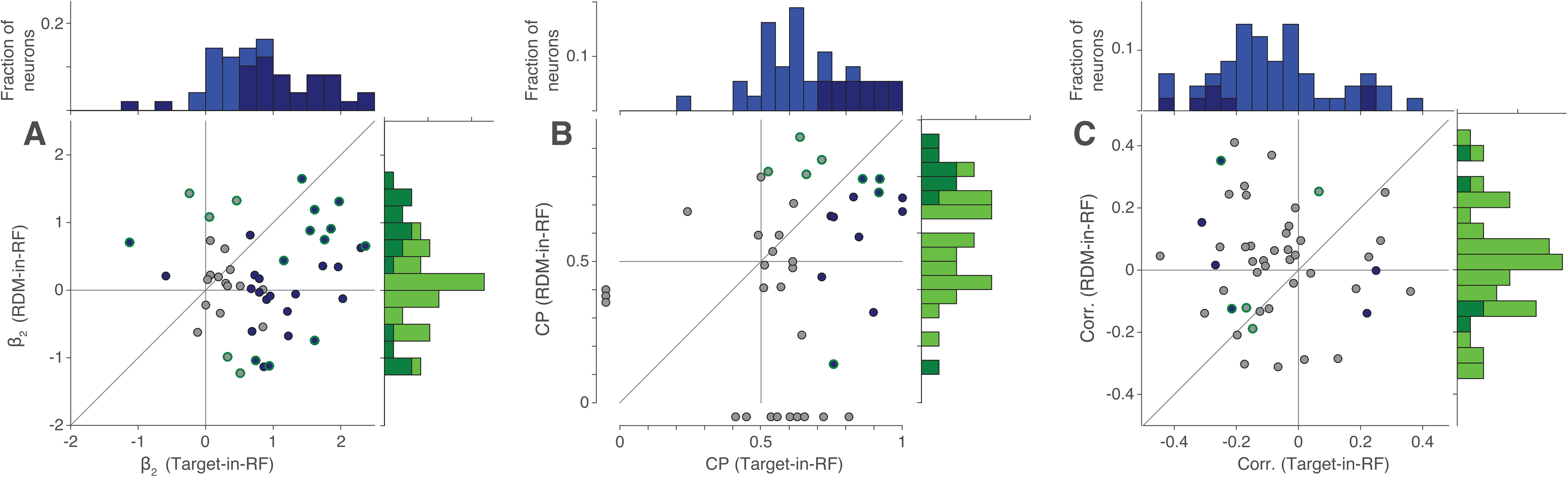
Leverage of neural activity on behavior. Scatter plot and histograms for the two stimulus configurations showing the distribution of *β2* term ***(A)*** of logistic regression (Equation 5), choice probability ***(B)*** and coefficient of correlation ***(C)*** between slope of response buildup and RT. Neurons for which the metric was significant are shown with a blue fill (significant in the Target-in-RF configuration) and/or a green border (significant in the RDM-in-RF configuration) in the scatter plots and as darker colors in the histograms. Data points in ***B*** outside the axes indicate neurons where choice probability could be determined for only one of the two configurations. One and three such data points are not shown in the scatter plots of ***A*** and ***C*** respectively.

We compared the strength of direction selectivity in our neural population to that reported in Fanini and Assad (2009), using their direction selectivity index (DI):

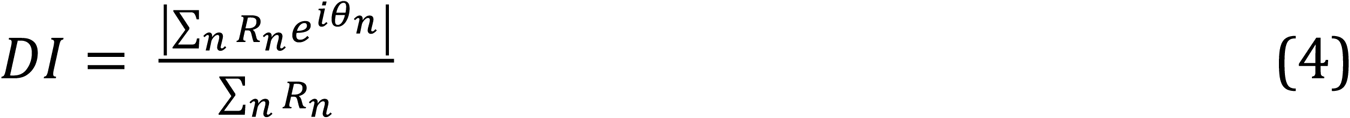

where *R_n_* is the mean response to *n^th^* direction *θn* in the time window 190 ms after RDM onset to 100 ms before saccade. DI was computed from responses to the 51.2% coherence motion trials in the two directions (*π* radians apart). We compared the distribution of the DI values in our population to those reported in Figure 3A of Fanini and Assad (2009), using a rank sum test (Figure 3E).

We used responses at the two strongest motion strengths (±51.2% coherence) to estimate the latency from motion onset to the time that direction selectivity was first apparent in a given neural population (Figure 3F). We averaged the responses in 40 ms bins on each trial at these coherences and derived receiver operating characteristics (ROC) from these response distributions at each time bin. The area under the ROC denotes the probability of the neuron responding more to the positive direction of motion. For each time bin, we applied a Wilcoxon rank sum test and estimated the response latency as the first of three successive bins that met statistical significance (p<0.05). We used a bootstrap procedure to estimate the distribution of latencies under the two task configurations. For each configuration, we resampled trials with replacement, matching the number of trials in the original data sets, and obtained a latency using the same procedure as on the actual data. We repeated this procedure 1000 times for each configuration. The medians of these distributions recapitulated the latency estimated from the data (180 and 190 ms for the Target-in-RF and RDM-in-RF respectively). We report the p-value of a rank sum test (2-tailed) using the bootstrap derived distributions to evaluate the null hypothesis that the latencies are the same for the two configurations. We obtained the same result by sampling neurons (instead of trials), with replacement.

We quantified the effect of motion strength on the rate of increase of neural response (‘buildup rate’) during the decision-making epoch as the slope of the response in the time window 180 to 380 ms after stimulus onset (Figure 3G). The start of the time window was chosen based on the latency of the direction selectivity of the responses. To exclude pre-saccadic activity, we discarded from each trial, the spikes occurring up to 100 ms before saccade onset. We computed by least squares method, the slope for each neuron at each coherence from the mean detrended response in 10 ms time bins in the aforementioned time window. We then tested whether these buildup rates scaled with coherence across the population in each stimulus configuration by fitting a linear model regressing these buildup rates against signed coherence. We confirmed that the trends shown in Figure 3G were preserved when the analysis was performed using weighted regression.

#### Leverage of neural activity on behavior: (Figure 4)

We measured the leverage of neural activity on the animal’s choice in two ways. First, we fit the monkey’s choices with logistic regression

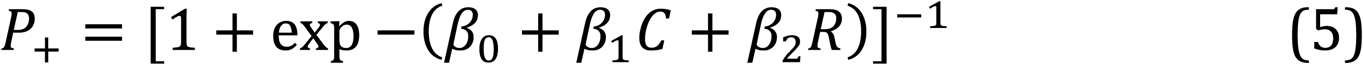

where *P+* is the probability of choosing the ‘positive’ direction target, *C* is signed coherence and *R* is the z-scored mean neural response in the time window 100 to 300 ms before saccade. If the variations in firing rate of the neurons have leverage over choice even when the effect of motion coherence is accounted for, then *β2* ≠ 0. We compared *β2* across configurations with a signed rank test on their absolute values. We also quantified the additional leverage of the neural responses on choice beyond that of the motion strength, by measuring the difference in the deviance of the full model and the model without the *R* term (*∧*). Comparisons of *∧* provided similar results to the comparisons of the *β2* term that are presented in the results.

Second, we quantified the trial-by-trial correlations between neuronal response and the animal’s choice in the 0% coherence trials by computing ‘choice probability’ (CP, Britten et al 1996). For each neuron, we computed the mean responses on the 0% coherence trials in a time window 100 to 300 ms preceding the saccade. The trials were separated into two groups based on the animal’s choice. We used the distributions of responses from the two groups to calculate the area under the ROC, termed the choice probability. We evaluated the null hypothesis that |CP-0.5|=0 using a permutation test. We permuted the union of responses from both groups and assigned them randomly to the two choices (matching the number of trials in each group) and computed the CP. By repeating this procedure 2000 times, we established the distribution of |CP-0.5| under *H0* and report the *p* value as the area to the right of the observed CP minus 0.5.

To evaluate whether the CPs from the two configurations were different, we first converted responses to z-scores (by neuron and configuration) and then combined the z-scores across neurons. We then computed two CPs, as above, for the two configurations. To evaluate the null hypothesis that the two CPs are equal, we performed another permutation test, this time preserving the association with choice but permuting the association with configuration. We obtained the distribution of the difference in CP (|ΔCP|) under *H0* from 2000 repetitions of the permutation procedure and report the p value as the area of this distribution that is greater than the observed |ΔCP| from the data.

We also quantified the correlation between the buildup rates and RT. We used trials in which the monkey chose the ‘positive’ direction target, including all such trials at 0% motion strength and only correct trials at positive motion strengths. For each trial, we computed the slope of the response between 180-420 ms after RDM onset (using 40 ms time bins) from the detrended responses. To remove the effect of coherence on RT, we standardized (i.e., z-scored) both the RTs and the buildup rates within each coherence and computed the correlation between them.

#### Variance and correlation analysis

To evaluate if the neuronal firing rates on individual trials during the decision-making epoch reflect a process of accumulation of noisy evidence, we analyzed the pattern of variance and autocorrelation of the responses (Churchland et al 2011, de Lafuente et al 2015). We were interested in the variance attributable to such an accumulation process. For the *i*^th^ time bin, this variance (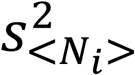) is the fraction of the total measured variance (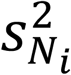) remaining after accounting for the point process variance (PPV), that is, the variance expected even if the underlying rates were constant. We refer to 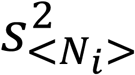, which is a variance of a conditional expectation of the counts, hence the variance of the underlying rate, simply as ‘variance’ in the main text. Assuming the PPV is proportional to the mean count,

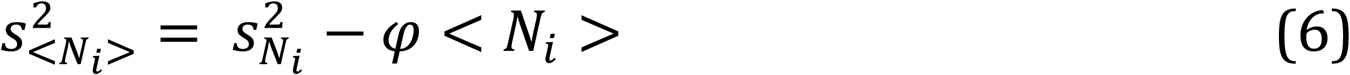

where *φ* is a constant that must be estimated.

Since our goal was to compare how well the firing rates conform to a diffusion process, we allowed *φ* to be a free parameter and fit it to obtain the best conformity to the autocorrelation pattern for a running sum of independent, identically distributed random numbers. Recall that the variance of the sum of *n* independent random samples of variance *σ^2^* is *nσ^2^*. If the sum is extended for another *m* samples, the variance is (*n*+*m*)*σ^2^*. The sum out to *n* shares a fraction of this variance: *n*/(*n*+*m*). This is the *R^2^*, and its square root is the correlation, *ρ*. So, for an unbounded diffusion process, the correlation between the *i*^th^ and *j*^th^ time steps is

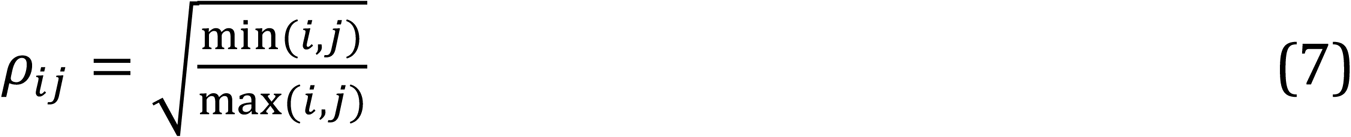

Note that for six time bins, the 6 by 6 correlation matrix contains 15 unique values of *ρi≠j*.

We characterized the variance and autocorrelation from six 60 ms time bins between 180-540 ms after stimulus onset, ignoring any time bins that extended to within 100 ms of the saccade. To pool data across neurons, we used the residuals for each trial as follows. The mean response of a trial in each time bin was subtracted from the mean of the responses from all the trials for that neuron for the same signed coherence in that time bin. We computed the covariance matrix from the residuals for the six time bins.

We used an initial guess for *φ* to calculate the variance attributable to the diffusion process (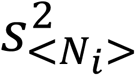, Equation 6) and substituted the raw variances for the diagonal of the covariance matrix. The correlation was derived from this covariance matrix by dividing each term by 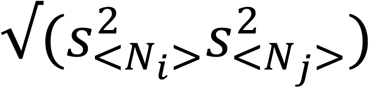. We used Nelder-Mead simplex method (MATLAB function ‘*fminsearch’*) to find the *φ* that minimized the sum of squares of the difference between the 15 z-transformed calculated correlation (*rij*) and the z-transformed theoretically predicted correlation (*ρij*). Note that the values of *φ* were not constrained to be the same in the Target-in-RF (*φ* = 0.42) and RDM-in-RF (*φ* = 0.39) configurations.

We report the variance 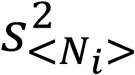) in Figure 5 using the fitted *φ* values and estimated the standard errors from a bootstrap. We evaluated the effect of time on the variance using least squares regression. We also performed these analyses over a range of plausible values of *φ* and confirmed that only the absolute values of the variances differed, whereas the shape of the variance function over time was unaffected. We similarly computed the variance and its standard error for time bins aligned to the onset of the saccade.

**FIGURE 5:**
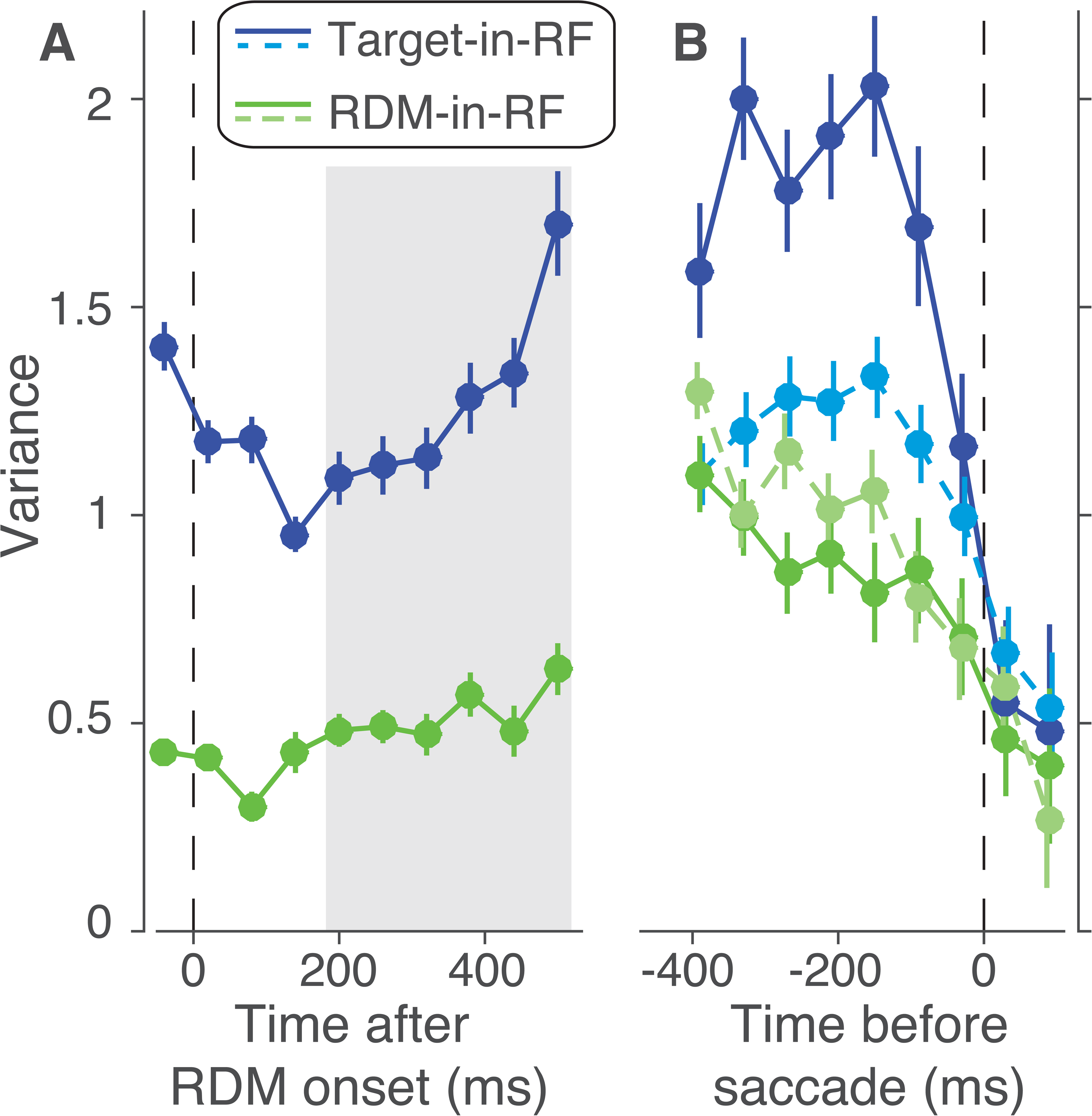
Variance of responses. The variance of neural responses aligned to the onset of RDM ***(A)*** or to the saccade ***(B)***. Total variance is computed in 60 ms bins and the point process variance subtracted from it (see Methods). In ***B***, solid lines are data from trials in which the animal chose the preferred target of the neuron and dashed lines are from trials with the opposite choice.

We used a combination of Monte Carlo methods and parametric statistical tests to analyze the decline in variance preceding the saccade. For trials in which the monkey chose the target in the RF, we compared the variance in the two time bins immediately preceding the saccade, using the bootstrap derived standard errors. We report a t-test. We made the same comparison for each of the other conditions: (1) unchosen Target-in-RF, (2) preferred direction choice with RDM-in-RF, and (3) non-preferred direction choice with RDM-in-RF. None were significant (p>0.05). We do not report these tests in the results and instead compare directly the estimates of variance decline in the four conditions. To do this, we computed the fractional difference in variance in the two time bins and estimated its standard error using the same bootstrap. We compared this difference statistic in the four conditions using ANOVA. We report the maximum *p* value for the comparison of the chosen Target-in-RF condition with the other three conditions, using Tukey’s test.

To quantify how well the measured correlation values conform to theoretical predictions, we formed a sum of square (SS) statistic from the 15 pairs of observed and theoretical correlations (after Fisher-z transformation, Figure 6D-E). We used a bootstrap procedure to estimate the distribution of this statistic by sampling with replacement from the data and following the steps above (100 iterations). We used a Kolmogorov-Smirnov test to determine the significance of the difference between the distribution of the SS statistics between the RDM-in-RF and the Target-in-RF configurations.

**FIGURE 6:**
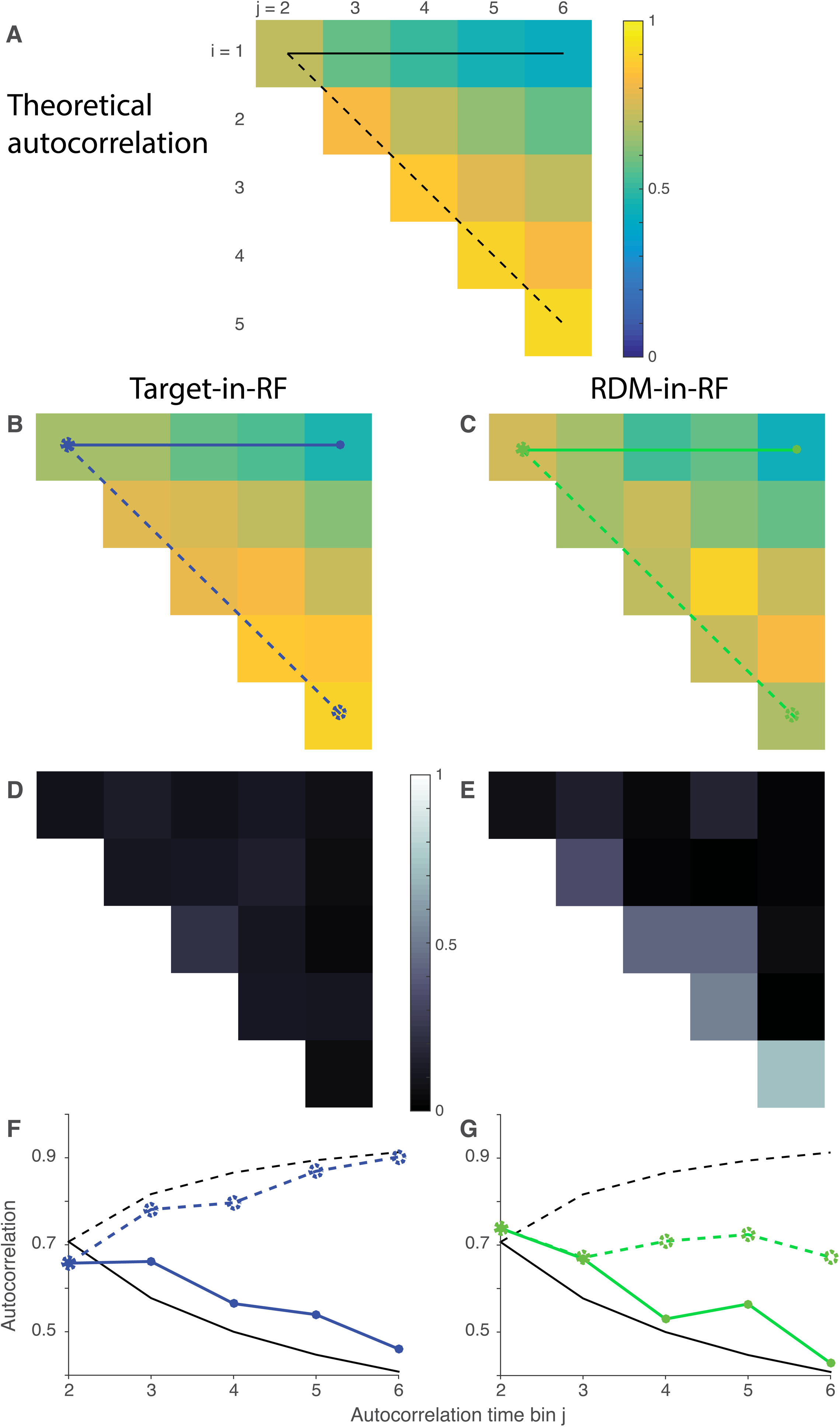
Autocorrelation of responses. ***A:*** Theoretical prediction of the autocorrelation matrix for six time bins (*ρi,j*) of a diffusion process. Only the 15 unique values (upper triangular matrix, *i<j*) are shown. ***B,C*:** Estimated autocorrelation for the neural responses in the two stimulus configurations. ***D,E:*** Deviation of *B,C* from the theoretical predictions shown in *A*. ***F,G:*** Comparison of correlation values in *A-C* between theory (black lines) and data (colored lines). Solid lines are correlation along the top row (between first and *j*^th^ time bins) and dashed lines along the first juxtadiagonal (correlation between *j*^th^ and its preceding time bins). Line style and color correspond to those in panels A-C.

### Model

We simulated the spike rates of three neural populations during the RDM epoch — one population with the RDM in their RF and two with targets in their RF. We devised two models that could account for direction selectivity seen in the RDM-in-RF population: (1) selectivity is inherited by means of divisive suppression from the Target-in-RF populations that are accumulating evidence (‘divisive suppression model’), and (2) selectivity arises from an evidence accumulation process transpiring in the RDM-in-RF population itself (‘parallel diffusion model’). Each model was implemented in two stages. In the first stage, our goal was to approximate the pattern of mean responses seen in the data. The models specify the predicted autocorrelation matrices for both neural populations. In the second stage, we compared the two models by assessing their capacity to explain the autocorrelation matrices derived from the neural data.

In the divisive suppression model (Figure 7A), the RDM-in-RF population was modeled as having an exponential rise in firing rate starting 50 ms after RDM onset and peaking at 130 ms (Figure 7C). The peak response varied from trial to trial, independent of RDM direction. The population then maintained the peak response through the end of the simulated epoch (540 ms after RDM onset). The two Target-in-RF populations were modeled as maintaining a steady response (*R*_0_) up to 180 ms after RDM onset and then following drift diffusion dynamics (Figure 7B). The responses *S* in the dynamic epoch evolved at each time step Δ*t* as

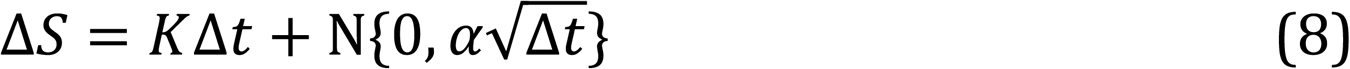

incorporating a deterministic drift component (*K*) and a diffusion component (N) — a Normally distributed random number with mean zero and standard deviation 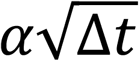. The drift component was positive for one target population (T1) and negative for the other (T2). The parameter K was chosen so that the drift rate in the T1 population of the model after implementation of divisive suppression (see below, Equation 9) matched the observed buildup of the neural response for the Target-in-RF neural population at the 25.6% coherence condition (solid line in Figure 7F). The parameter *α* was chosen such that the slope of the variance, after incorporation of suppression, mimicked that seen in data (blue curve in Figure 5A). See Table 2 for values of model parameters.

**FIGURE 7:**
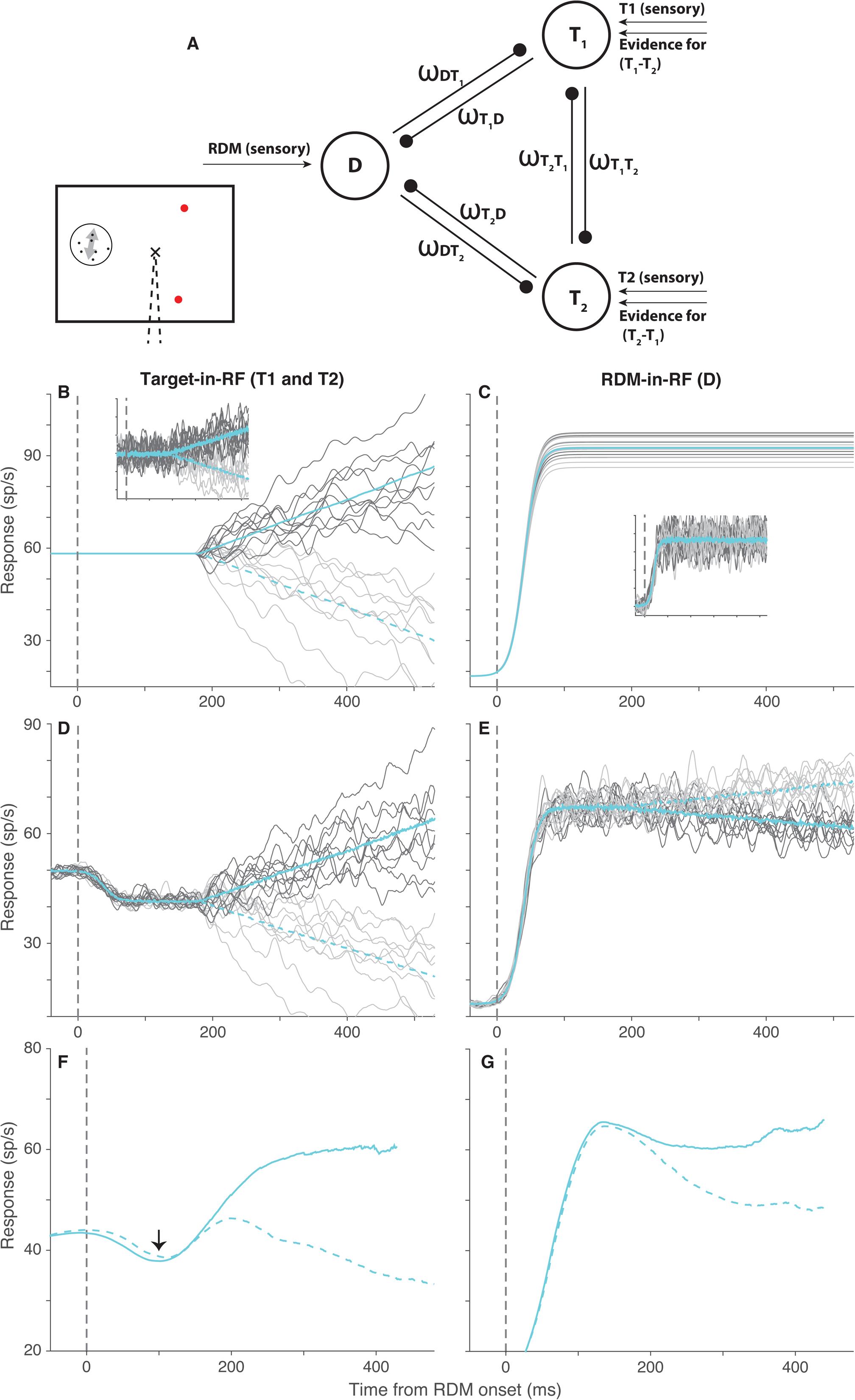
Divisive suppression model. ***A:*** Schematic of the three populations simulated in the model – one population representing the RDM (D) and two representing the targets (T1 and T2). The *ω* terms denote the suppressive influence of each population on the other two. ***B:*** Average response of simulated T1 (solid cyan) and T2 (dashed cyan) populations across trials in which the direction of motion supported T1. Dark and light gray traces show responses to 10 example trials for the two populations. ***C:*** The mean and example trial responses of the D population to the two directions of motion. Dark and light gray indicate motion towards T1 and T2, respectively. Solid and dashed cyan lines denote the corresponding average response traces, but they overlap, as the two populations do not distinguish between directions of motion. Insets in B and C show the noisy versions of the corresponding responses that furnish the divisive suppression. ***D,E:*** The responses of the three populations after implementation of divisive suppression. Color scheme is the same as in panels *B* and *C*. The simulated responses in *B-E* are smoothed with a 10 ms boxcar filter. ***F,G*:** The average responses of the recorded neural population to the 25.6% motion strength stimulus in the Target-in-RF and RDM-in-RF configurations that our simulations approximated. These traces are the same as the cyan traces in Figure 3A & C.

**Table 1:** Bounded diffusion model best fit parameter values (±SE)

| Parameter | Monkey N<br>(Target-in-RF) | Monkey N<br>(RDM-in-RF) | Monkey B<br>(Target-in-RF) | Monkey B<br>(RDM-in-RF) |
| --- | --- | --- | --- | --- |
| $\kappa$ | $16.05 \pm 0.38$ | $13.86 \pm 0.44$ | $9.66 \pm 0.39$ | $12.00 \pm 0.70$ |
| $B_0$ | $0.72 \pm 0.02$ | $0.78 \pm 0.02$ | $0.52 \pm 0.02$ | $0.47 \pm 0.04$ |
| $B_{del}$ | $0.01 \pm 0.00$ | $0.00 \pm 0.01$ | $0.02 \pm 0.01$ | $0.02 \pm 0.01$ |
| $B_2$ | $0.67 \pm 0.09$ | $0.97 \pm 0.08$ | $1.16 \pm 0.21$ | $1.26 \pm 0.38$ |
| $t_{nd1}$ | $0.34 \pm 0.01$ | $0.29 \pm 0.01$ | $0.41 \pm 0.01$ | $0.45 \pm 0.01$ |
| $\sigma_{tnd1}$ | $0.13 \pm 0.00$ | $0.11 \pm 0.00$ | $0.07 \pm 0.00$ | $0.08 \pm 0.00$ |
| $t_{nd2}$ | $0.38 \pm 0.01$ | $0.32 \pm 0.01$ | $0.47 \pm 0.01$ | $0.43 \pm 0.01$ |
| $\sigma_{tnd2}$ | $0.12 \pm 0.00$ | $0.12 \pm 0.00$ | $0.07 \pm 0.00$ | $0.06 \pm 0.00$ |
| $C_0$ | $0.00 \pm 0.00$ | $0.00 \pm 0.00$ | $-0.02 \pm 0.00$ | $0.02 \pm 0.01$ |

**Table 2:** Parameter values for simulations.

| Parameter | Divisive suppression model | Parallel diffusion model |
| --- | --- | --- |
| $K(T)$ | 80.4 | 52.8 |
| $\alpha(T)$ | 29.8 | 23.7 |
| $K(D)$ | N/A | 25.0 |
| $\alpha(D)$ | N/A | 9.6 |
| $\omega_{T2T1} = \omega_{T1T2}$ | $2 \times 10^{-3}$ | N/A |
| $\omega_{DT1} = \omega_{DT2}$ | $4 \times 10^{-3}$ | N/A |
| $\omega_{T1D}$ | $6 \times 10^{-3}$ | N/A |
| $\omega_{T2D}$ | $1 \times 10^{-3}$ | N/A |
| $\phi_{RDM}$ | 0.38 | 0.39 |
| $\phi_{Tar}$ | 0.43 | 0.43 |
| $V_{RDM}$ | 4.17 | 0 |

We simulated 10,000 trials and implemented divisive suppression between the three populations of the form

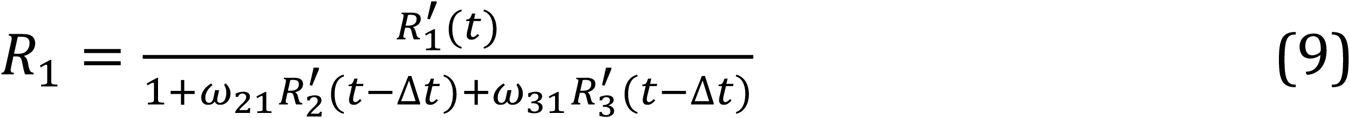

where *R’* and *R* denote the unsuppressed and suppressed responses, respectively, of the population indicated by the subscript, and *ω_ij_* is the weight of the influence of the *i*^th^ population on the *j*^th^. The suppressed responses at each time point (*t*) was computed based on the unsuppressed responses in the time window preceding it by Δ*t* = 10ms.

We first estimated the suppression of two target populations on each other (*ω*_vCvF_ and *ω*_vFvC_) from the peak and steady state responses of the neurons to the appearance of a target in their RF. We then estimated the weight of suppressive influence of the RDM-in-RF population on the Target-in-RF populations (*ω_DTx_*, *X* ∈ {1,2}) using the firing rates at the trough of the response dip following the onset of RDM (arrow in Figure 7F). The influences of the two Target-in-RF populations on the RDM-in-RF population *ω*_vxw_ were adjusted around *ω*_wvx_ to mimic the observed separation in mean responses of the RDM-in-RF population to the two directions of motion. Such asymmetry of the influence of the two Target-in-RF populations might arise from the different spatial relationship they might have with the RDM-in-RF population. Similar asymmetries are likely for the other pairs of *ω* too, but we set them to be equal here to simplify the model. We used the weights of suppression to estimate the underlying unsuppressed mean responses of each of the populations (Figure 7B-C).

In the parallel diffusion model, we implemented drift diffusion dynamics in the RDM-in-RF population as well as in the Target-in-RF population, and the populations had no suppressive interactions (Figure 9). The drift component in the RDM-in-RF population (*K* in Equation 8) was set to mimic the observed separation of responses to the two directions of motion in the data (Figure 7G). The scaling factor for the variance of the diffusion component (*α* in Equation 8) was adjusted to mimic the observed slope of the variance of the responses in the RDM-in-RF configuration (green curve in Figure 5A). Because of the absence of divisive interactions in this model, *K* and *α* for the Target-in-RF populations were recomputed to bring them in agreement with the data (Table 2).

**FIGURE 8:**
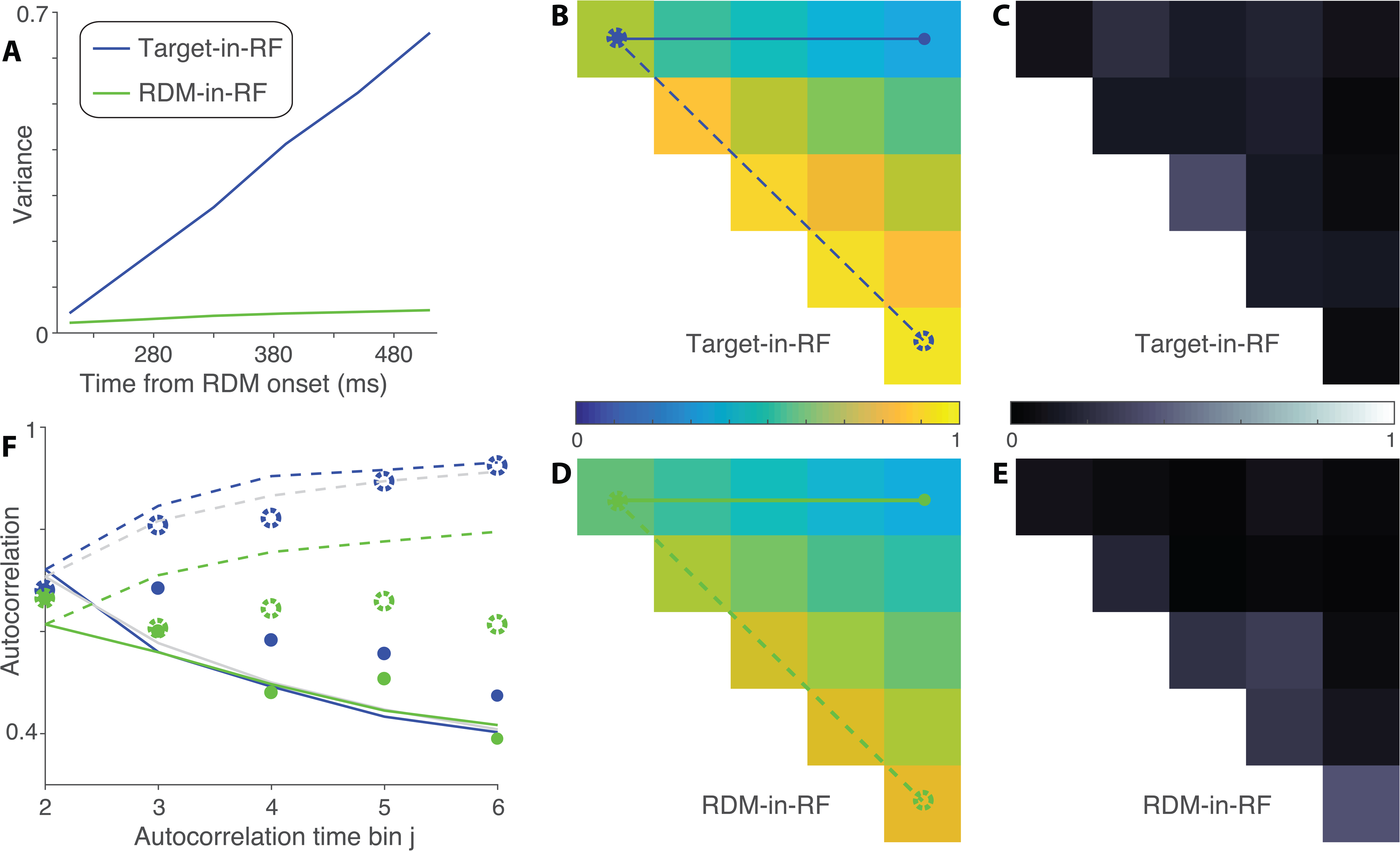
Variance and correlation of the simulated responses. ***A:*** Variance as a function of time in two of the simulated suppressed populations (D and T1 for trials with motion supporting T1 choice). ***B:*** Autocorrelation in the simulated suppressed Target-in-RF population T1. Conventions as in Figure 6B. ***C:*** Deviation of the autocorrelation in the model from the autocorrelation estimated from the data in the Target-in-RF configuration. ***D,E:*** Same as B and C for the RDM-in-RF population. ***F:*** Comparison of correlation values along the top row (solid lines) and first juxtadiagonal (dashed lines) between the model (see panels *B* & *D*) and the data. Circles show the correlation estimated from data. Filled circles correspond to the values along the top row and open circles to the values along the juxtadiagonal. Gray lines show the correlation expected from a diffusion process.

**FIGURE 9:**
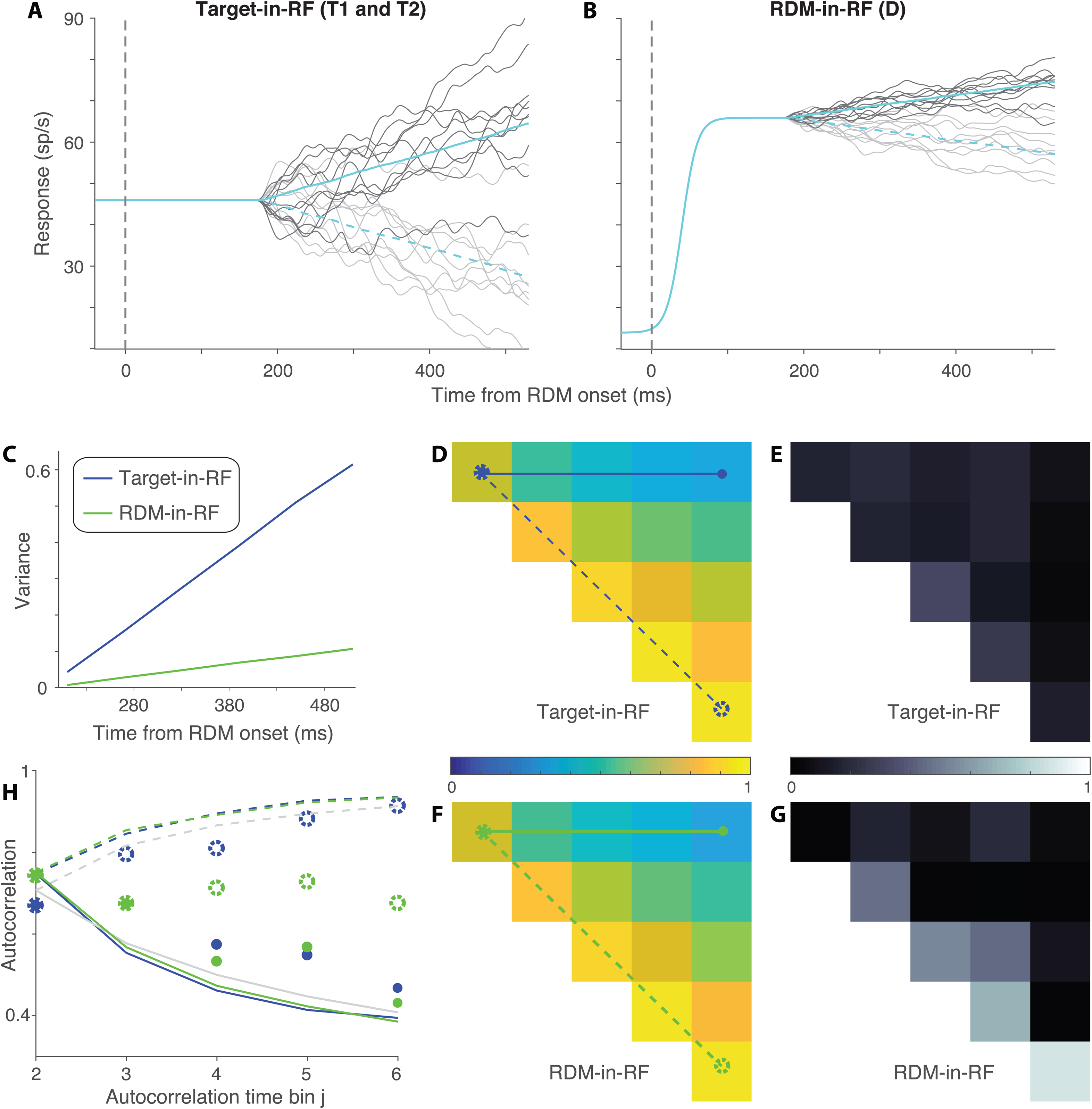
Alternative model with the RDM-in-RF population showing drift-diffusion dynamics. This model assumes that there is no interaction between the RDM-in-RF neurons and the Target-in-RF neurons. ***A-B:*** Responses of model neurons. Respectively similar to Figure 7B-C. ***C-H:*** Variance and autocorrelation in the model and the data. Conventions as in Figure 8A-F respectively.

Up to here, all parameters were established from the neural data, allowing both models to approximate the mean responses in the data. To compare how well the two models can account for the pattern of autocorrelation in the data, we needed to consider other possible sources of variance and autocorrelation. In both models, the variance of the non-directional sensory response of the RDM-in-RF populations was incorporated as a free parameter *V_RDM_*. This parameter was constrained to not exceed the variance observed at the peak of the sensory neural response in the RDM-in-RF configuration. For the divisive suppression model, our hypothesis is that the noisiness of the suppression causes the autocorrelation pattern of the RDM-in-RF population to deviate from theoretical predictions. We instantiated this noisy process by corrupting the interaction signals so that they were not perfect replicas of the responses of the three populations in the model (Insets in Figure 7B, C). This noise term was proportional to the square root of the response. We set the scaling term *γ*=5 to represent a modest amount of noise (*R*^2^= 0.81 for the diffusion paths and their corrupted versions).

We attempted to achieve the best possible fit to the 30 correlations observed in the data in the two configurations (15 unique values each for the Target-in-RF and RDM-in-RF configuration) under each of the models. The models give rise to predicted correlations in the Target-in-RF and RDM-in-RF populations (varying with the free parameter *V_RDM_*). As above, we allow for uncertainty in the PPV in the data (*φ* in Equation 6). So we compute the correlations in the neural data with two additional degrees of freedom (parameters, *φ_RDM_* and *φ_Tar_* for the RDM-in-RF and Target-in-RF configurations, respectively). We estimated the set of parameters that maximized the log likelihood (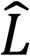) of the 30 correlations in the data (Fisher z-transformed) under the model predictions. It was not possible to fit *γ* and *φ_RDM_* simultaneously without imposing additional constraints (e.g., *φ_RDM_*=*φ_Tar_*). Instead, we fixed *γ* to establish a modest perturbation of the interaction signals, as noted above. This is the model illustrated in Figures 7-8 (parameters in Table 2). We compared models using the difference in Bayesian Information Criterion (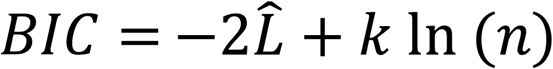, where *k* is the number of free parameters and *n* is the number of data points). We explored a range of *γ*, to confirm that the suppression model is favored even with subtle noise perturbation (e.g., ΔBIC>100 for *γ*=1, *R*^2^= 0.99). BICs were calculated by conservatively assuming 4 degrees of freedom (d.f.) for the divisive suppression model {*φ_RDM_, φ_Tar_, γ*, *V_RDM_*} and just two d.f. for the parallel diffusion model {*φ_RDM_, φ_Tar_*} because *γ* should be regarded as a free parameter and the best fit of the parallel diffusion model assigns *V*RDM ≈ 0. We also fit to a model with *γ* as a free parameter under the constraint *φ_RDM_*=*φ_Tar_*. This implementation also favors the suppressive interaction model (ΔBIC>6×10^3^; best fitting *γ*=8.2). Note that the implementation of *V_RDM_* introduces autocorrelation of the rate that spans the duration of the analysis epoch (360 ms). Parametrization of the sensory responses with exponentially decreasing autocorrelation did not provide a significantly better fit to the data in either model.

## RESULTS

We recorded from 49 well isolated single neurons in area LIP from two monkeys (28 neurons from monkey N and 21 neurons from monkey B) as they decided the net direction of a noisy random-dot motion (RDM) stimulus. On each trial, two choice targets indicated the two directions to be discriminated (e.g., up vs. down). The monkeys reported their decision by making a saccade to the choice target along the perceived direction of motion. They were free to indicate their decision whenever ready, thus providing a measure of reaction time (RT). The monkeys performed the task with the RDM and the targets arranged in two configurations (Figure 1). In the ‘*Target-in-RF*’ configuration, one of the choice targets was placed in the response field (RF) of the neuron under study. In the ‘*RDM-in-RF’* configuration, the RDM was placed in the RF. In this way, we obtained data from the same LIP neuron when it belonged either to the pool representing the RDM stimulus or to one of the two pools representing the choice targets.

We first establish that the animals integrate motion information over 100s of ms to make their choices in both task configurations. This prolonged deliberation time offers a window in which to interrogate how the neural responses relate to the process of decision formation. We show that the firing rates of neurons represent the state of the accumulated evidence only when the neurons belong to a pool representing the targets.

### Behavior in the two task configurations

The behavior of both monkeys exhibited an orderly dependence on the strength of the RDM in both task configurations. They took longer to report their decision when the motion strength was weaker (Figure 2, A-D), and their decisions were less accurate (Figure 2, E-H). The systematic relationship between reaction time (RT) and accuracy is well described by the accumulation of noisy evidence to a threshold, which determines both the time it takes to make a decision and which alternative the monkey chooses (Gold & Shadlen 2002, Smith & Ratcliff 2004). We support this assertion by fitting the RTs to a bounded evidence accumulation model and then using the fitted parameters to predict the choices (Kang et al 2017, Shadlen & Kiani 2013). Specifically, the curves in the top row of Figure 2 are fits to a parsimonious symmetrically bounded drift diffusion model, which uses four parameters to account for the effect of motion strength on the mean RT for correct choices (Equation 1; see Methods). Two of the parameters—the bound height, ±B, and the sensitivity coefficient, *κ*—establish predictions for the proportion of choices as a function of motion strength (Equation 2). The dashed curves in the lower panels of Figure 2 depict these predictions. They are only slightly worse than logistic fits to the choice data themselves (gray curves), which are unconstrained by RT. To quantify the “goodness of prediction”, we compared the model predictions to those obtained from random perturbations of the mean RTs which preserve their orderly dependence on motion strength. Small perturbations of the RT (mean 7.5%, range 1-12% or equivalently, mean 48 ms, range 7-73 ms) are sufficient to produce substantially poorer predictions (p<0.01). The fidelity of the predictions supports the assertion that the choices result from the same process of bounded evidence accumulation that explains the decision times. Importantly, this conclusion holds for both stimulus configurations.

From this exercise we conclude that the decision times (i.e., RT minus the non-decision time) estimated from diffusion model fits can be used to identify an epoch in which noisy evidence was integrated to make the decision. To obtain more refined estimates of the integration times for the different task configurations, we fit a more elaborate bounded diffusion model (Figure 2-Extended data Figure 1, see Methods for details and Table 1 for fit parameters). The small differences in reaction times between the two configurations for Monkey N was accounted for by the non-decision time parameter. For Monkey B, a combination of increased sensitivity and decreased bound height contributed to the faster RTs in the RDM-in-RF configuration. Importantly, the fits established that both monkeys integrated evidence over hundreds of ms in each configuration.

### LIP neuronal responses in the two task configurations

Neurons in area LIP can exhibit sensory-, memory- and saccade-related responses (Barash et al 1991a, Gnadt & Andersen 1988). For example, in a task where a monkey must remember a visually cued location and make a delayed saccade to it, LIP neurons can show (1) a short latency response to the visual cue if it appears in the RF, (2) a persistently elevated response during the delay period and (3) a burst of activity preceding a saccade to the remembered location. Not all LIP neurons exhibit all three types of responses. Since our goal was to compare the decision related activity in the same neurons when they belonged to the pool representing the sensory information and when they belonged to the pool involved in planning the motor action, we recorded from neurons that responded to visual stimuli in their RF and also showed persistent activity in association with saccadic motor planning. Each of our neurons increased their responses above baseline to the appearance of a visual stimulus in their RF (responses after RDM onset: median 5 SD above baseline, interquartile range [IQR]: 2.7 to 7.7). The strength of this sensory response was comparable to the highest responses observed during the delay period (median 4.3 SD above baseline, IQR: 2.3 to 9.2, p=0.49, Kolmogorov-Smirnov test).

During the direction discrimination epoch, the pattern of activity of the recorded neurons varied according to which pool they belonged to. When the neurons belonged to a pool with one of the targets in the RF, the responses largely recapitulated observations from earlier reports (e.g. Churchland et al 2008, Roitman & Shadlen 2002). Figure 3 shows the average population response of all neurons in the Target-in-RF configuration, aligned to either the onset of RDM (Figure 3A) or to the saccade (Figure 3B). The response was elevated before the onset of the RDM reflecting the presence of a choice target in the RF of the neurons. Following motion onset, there was a stereotyped dip in activity before the responses began to separate by motion strength. The evolution, beginning ∼180 ms after stimulus onset, is best appreciated in the de-trended responses (Figure 3A, inset). These features and those next described were evident in both of the monkeys, shown individually in Figure 3-Extended data Figure 1 and 2.

The same neurons also exhibited differential responses to the two directions of motion being discriminated when they belonged to the pool representing the RDM. To combine responses across the population in this task configuration, we identified the preferred direction of motion for each neuron as the one that elicited the greater response. Figure 3C-D shows the responses of the population averaged after sorting by each neuron’s preferred direction. After an initial rise in activity due to the appearance of the RDM in the RF, the responses exhibited a direction dependent separation. Such modulation of LIP neuronal responses by motion direction has been previously reported in naïve monkeys (Fanini & Assad 2009). However, the direction dependent modulation was slightly stronger in our neural population (median direction selectivity index: 0.11 and 0.09, respectively for our neurons and those reported in Fanini & Assad; p=0.06 rank-sum test; see Figure 3E). Note that, our neural population displays this degree of direction selectivity at a lower motion strength (51.2% coherence) than that used by Fanini & Assad (100% coherence). This result is consistent with previous reports of stronger directional selectivity in LIP neurons of monkeys trained on tasks that rely on direction discrimination (Sarma et al 2015).

We quantified the time course of the evolution of direction selectivity at the highest motion strength (Figure 3F) using an ROC metric (see Methods). The responses to the two motion directions were significantly different starting 190 ms after the onset of dot stimulus (p<0.05 on Wilcoxon rank sum test). This is much later than the ∼50 ms latency of direction selectivity observed in naïve monkeys (Fanini & Assad 2009). This is also later than the ∼100 ms latency for direction category selectivity reported in monkeys trained to categorize sets of motion directions (Swaminathan & Freedman 2012). As discussed below, the long latency in our neuronal pool may be an indication that the directional responses we observed in the RDM-in-RF configuration arise through a different mechanism than the direction- and category-selective responses previously reported in LIP.

The latency in the RDM-in-RF configuration lagged the direction selectivity seen in the same neurons in the Target-in-RF configuration (180 ms, p<10^−3^, bootstrap analysis). However, the similarity of the latencies suggests that the RDM-in-RF population might also reflect the formation of the decision, as the Target-in-RF population has been shown to do (Churchland et al 2008, Roitman & Shadlen 2002). Consistent with this possibility, the rise and decline of neural activity depends on the strength of the RDM (Figure 3C, inset), albeit with a smaller dynamic range compared to responses in the Target-in-RF configuration. Note that in this configuration, directions are sorted based on the preferred direction of each neuron. The coherence dependent ordering of responses could have been accentuated by this *post hoc* procedure. To quantify this coherence dependence, for each neuron and motion strength, we estimated the slope of the responses (buildup rate) in a 200 ms epoch beginning at the time of response separation as identified in the preceding analysis. We then characterized the relationship between motion strength and buildup rates separately for the preferred and non-preferred directions of motion (Figure 3G). The buildup rates of neurons in the Target-in-RF configuration showed a linear dependence on motion strength both when the motion direction was towards the RF (1.5±0.2 spikes per s^2^ per 1% coherence, p<10^−9^) and when the motion was away from the RF (−1.2±0.2, p<10^−5^). A similar trend was observed in the RDM-in-RF configuration. However, this relationship was significant only for the non-preferred direction of motion (−0.7±0.2 spikes per s^2^ per 1% coherence, p<0.002). For the preferred direction, the build-up rates increased with coherence but not significantly so (0.6±0.4 spikes per s^2^ per 1% coherence, p=0.13). In both configurations, these trends were preserved even when the highest motion strength trials were excluded. Thus, neuronal pools in LIP representing the saccade targets and the RDM both differentiate the discriminanda during an epoch coinciding with decision formation. The build-up of neural activity depended on the strength of the stimulus in both populations, but this dependence was weaker when the RDM was in the RF.

We also compared the responses at the end of the decision process for the two task configurations (Figure 3B & D). When the monkey chose the target in the neuron’s RF, the responses appear to coalesce to a common firing rate just before the saccade, irrespective of motion strength (Figure 3B, solid curves), as shown previously (Churchland et al 2008, Roitman & Shadlen 2002). This pattern is thought to reflect a threshold level detected by another circuit to terminate the decision (Hanes & Schall 1996, Hanks et al 2014, Mazurek et al 2003). When the same neurons contained the RDM in their RF, the responses to the different coherences remained separated until the saccade, and this held for either choice (Figure 3D). This was also the case when the RF contained the unchosen target (Figure 3B, dashed curves). Thus, only the responses of the pool representing the target chosen by the animal contains a possible neural signature of decision termination. In the ensuing sections, we support this qualitative observation with other lines of evidence that show that this pool alone signals decision termination and the time taken to reach it.

### Correlation between neural responses and behavior

We examined whether the neural responses in the two stimulus configurations were predictive of the monkey’s decisions. Specifically, we asked if the trial to trial variation in the responses correlates with the trial to trial variation in the monkey’s choice behavior. To test this for each neuron, we counted the spikes in a 200 ms long epoch ending 100 ms before saccade initiation on each trial and incorporated this count in a logistic regression model of choice (GLM; see Methods). To facilitate comparison across the two stimulus configurations, we standardized the responses across trials of each configuration. We included the strength and direction of the presented stimulus as confounders, thus asking whether the variation in neural response tells us more about the upcoming choice than can be ascertained from the stimulus itself. This was indeed the case for 61.2% of cells in the Target-in-RF configuration and for 35.4% of cells in the RDM-in-RF configuration (30 of 49 and 17 of 48 cells respectively; Equation 5, *H0*: *β2* = 0; p<0.05; Figure 4A). The leverage of the neural activity on choice was significantly stronger in the Target-in-RF configuration (p=0.005, signed rank test).

In a complementary analysis, we assessed whether the neural responses on ambiguous trials (0% motion coherence) differed according to the eventual choice of the animal. We computed choice probability (Britten et al 1996), a nonparametric statistic that quantifies the overlap between the distributions of responses of the neuron accompanying the two choices (see Methods). A choice probability of 0.5 indicates that the two distributions are completely overlapping and therefore uninformative about the ensuing choice. At the single neuron level, choice probability of 32.4% and 25.8% of the neurons was significantly different from 0.5 in the Target-in-RF and RDM-in-RF configurations, respectively (12 of 37 and 8 of 31 cells with at least 10 trials at 0% coherence respectively, p<0.05, permutation test). In both stimulus configurations, the mean choice probability of the neuronal population was significantly greater than 0.5 (Figure 4B, population mean ± SEM of 0.66±0.03 and 0.59±0.04 for Target-in-RF and RDM-in-RF respectively, p<10^−5^ and p<0.02 on t-test). For comparison between the two configurations, we calculated ‘grand’ choice probability from standardized responses of all neurons on the 0% coherence trials (see Methods, Britten et al 1996). This choice probability was significantly stronger in the Target-in-RF configuration (0.65 vs. 0.56, p<10^−3^, permutation test). From the analyses of choice probability and firing rate leverage on choice (Figure 4A-B) we adduce that LIP neurons responsive to both the RDM and the choice targets are informative about the choice, but it is the latter set of neurons (Target-in-RF) that covary more strongly with choice.

Finally, since the neurons exhibit time dependent changes in their activity in both stimulus configurations, we asked whether the variation of the buildup rates were predictive of the variation in the RTs on a trial-by-trial basis. We used the trials in which the monkey chose the target in the RF or the target consistent with the direction of motion preferred by the neuron (RDM-in-RF). For a majority of neurons recorded in the Target-in-RF configuration (36 of 49), the reaction times were inversely correlated with the slope of the neural responses (population mean: −0.08, p<0.01). In the RDM-in-RF configuration, the correlation was not significantly different from 0 (mean: 0.03, p>0.33) (Figure 4C) and significantly weaker than the correlations seen in the Target-in-RF configuration (p<0.01, Kolmogorov-Smirnov test). This comparison suggests that only the pool of neurons that contain the chosen target in their RF carries information about the time the animal will take to report its decision.

### Signatures of noisy evidence accumulation in the response variance

We also wished to ascertain whether the responses on single trials conform to the expectations of noisy evidence accumulation. If so, the variance of the firing rates across trials should increase linearly as a function of time (i.e., the number of samples accumulated). Also, the autocorrelation between firing rates at different times within a trial should conform to the pattern associated with the cumulative sum of random numbers. Such correlation should decay as a function of separation in time from the first sample and increase for equidistant samples as a function of time from the onset of accumulation (see Methods). We used the method developed by de Lafuente et al (2015) (based on Churchland et al (2011)) to estimate these quantities.

The variance and autocorrelation patterns varied markedly based on whether the neurons contained the target or the RDM in their RF. In the Target-in-RF configuration, the variance increased linearly with time during the same epoch that the mean firing rates seemed to reflect the integration of evidence (Figure 5A, shaded region). In the RDM-in-RF configuration, the rise in variance was significantly weaker (p<10^−10^, bootstrap analysis). Also, the observed autocorrelation matrix for the responses in the Target-in-RF configuration (Figure 6B,D,F) resembled the theoretical prediction (*R*^2^ = 0.88). In contrast, the pattern of autocorrelations (Figure 6C,E,G) for the responses in the RDM-in-RF configuration diverged markedly from the predicted pattern (*R*^2^ = 0.2). A bootstrap analysis confirmed that the difference in *R*^2^ values between the two configurations was statistically reliable (p<10^−10^; see Methods). Later, we show that the deviation of the autocorrelation pattern from theoretical prediction cannot be attributed to a muted drift diffusion process unfolding on the background of a strong non-directional sensory response (Figure 9).

The variance of the neural response also affords a more refined examination of the mechanism of decision termination. The firing rate averages in Figure 3B suggest the possibility that decisions terminate when the firing rate of the neurons with the chosen target in their RF reach a threshold. A more stringent test of a threshold is that even for the same motion strength, the variance of the neural response should approach a minimum just before the saccade. Indeed, we observed a precipitous decline in the variance in the ∼100 ms preceding the saccade for the neuronal pool with the chosen target in the RF (Figure 5B, solid blue line). The variance in the time bin preceding the saccade was significantly lower than the variance in its prior time bin (p<0.01, t-test). This decline in variance was more precipitous than that seen for the other three conditions shown in Figure 5B (ANOVA, p<0.03, see Methods).

Together, the analyses of time dependent variance and autocorrelation reveal that neurons in the Target-in-RF configuration exhibit firing rate patterns consistent with a process that represents the running sum of noisy samples of evidence to a criterion level. The analyses complement the observations made earlier on the mean firing rates by demonstrating conformance with the second order statistics of diffusion to a bound. These features were less apparent when the same neurons were studied in the RDM-in-RF configuration. This neural population does not appear to represent the accumulation of the noisy evidence that supports the monkey’s decisions. They reflect the direction of motion during the time course of decision formation but not the state of the accumulated evidence that can be used to terminate the decision process. We next consider a possible account of their pattern of activity.

### A model of interaction between populations

How could the responses of neurons with the RDM in their RF correlate with the decision outcome without representing the process of evidence accumulation? One possibility is that the weaker decision-related signals observed in the population with the RDM in their RF are inherited from the populations that have the choice targets in their RF and are involved in the accumulation process. It has been shown that responses of LIP neurons to visual stimuli are suppressed by concurrently presented visual stimuli when they are well outside the RF (Balan et al 2008, Churchland et al 2008), even by as much as 50° visual angle (Falkner et al 2010, Louie et al 2011). An asymmetrical influence of the two Target-in-RF populations could lead to the appearance of direction selectivity and a correlation with the animal’s choices in the RDM-in-RF population. Moreover, the noise added through this additional step could explain the divergence of the variance and autocorrelation of the RDM-in-RF population from the theoretical predictions of a diffusion process. Additionally, such an extra step could account for the timing of direction selectivity in the RDM-in-RF population, which lags slightly behind that of the Target-in-RF population.

To evaluate the plausibility of this idea, we simulated the responses of three neural populations—one representing the motion stimulus and two representing the choice targets—during the motion viewing epoch (Figure 7A). In the model, the RDM-in-RF population receives direct excitation from the visual representation of the dynamic random dots. This direct excitation furnishes a constant firing rate that varies from trial to trial, but importantly, is not direction selective (Figure 7C). The two Target-in-RF populations start off at a steady firing rate, simulating the steady state sensory response to the target already present in the RF. The responses then follow drift-diffusion dynamics starting at 180 ms, simulating evidence accumulation. The drift rate was set to be directly or inversely proportional to motion coherence for the populations representing the correct and incorrect targets, respectively (Figure 7B).

The three populations interact through divisive suppression (Carandini & Heeger 2011, Louie et al 2011, Sceniak et al 2001) at each time point, parameterized by the ω terms in Equation 9 (Methods). We set these parameters to approximate the observed neural responses to the 25.6% motion strength RDM (illustrated in Figure 7F-G). We assumed that the early dip in the response of the Target-in-RF neurons (arrow, Figure 7F) was caused by suppression from the neurons activated by the appearance of the RDM (ωDT1=ωDT2). The suppression between the two Target-in-RF pools (ωT1T2=ωT2T1) was estimated from the onset and steady state responses after the appearance of the target in the RF. Suppression of the RDM-in-RF pool from the Target-in-RF pools (ωT1D and ωT2D) were adjusted around ωDT to approximate the separation in firing rate traces shown in Figure 7G (see Methods). Such asymmetric influence of the two Target-in-RF populations might arise from differences in their spatial relationship (neuronal connectivity) with the RDM-in-RF population. These adjustments were sufficient to mimic the observed mean responses of the neural population in our simulations (Figure 7D-E). In addition, we assumed that the suppressive interaction signals were corrupted by a small amount of noise (see Methods). Importantly, according to the model, the direction selectivity of the RDM-in-RF population is derived solely from the suppressive inputs from the Target-in-RF populations.

This simple model reproduced the main features of our results (Figure 8). After the implementation of suppression, the Target-in-RF population retained the time course of the variance and the pattern of autocorrelation expected of a diffusion process. Notably, the variance and autocorrelation in the RDM-in-RF population also conformed to the patterns in the neural data: (*i*) the attenuated increase in variance as a function of time and (*ii*) the divergence in the pattern of autocorrelation from the theoretical prediction of diffusion. We also considered an alternative model in which the RDM-in-RF population itself represents an attenuated evidence accumulation signal in parallel with the Target-in-RF populations (Figure 9). To do this, we removed the lateral interactions and implemented the accumulation identically to the Target-in-RF population, but matching the observed firing rate dynamics and variance in the RDM-in-RF data (displayed in Figures 7G and 5A, respectively). This model was significantly worse in accounting for the pattern of autocorrelation observed in the data (ΔBIC > 5×10^3^). We thus favor the model with divisive suppression, which accounts for the presence of choice related activity in the RDM-in-RF population and the absence of clear signs of noisy evidence accumulation.

## DISCUSSION

We compared decision related activity in the sensory and motor-planning responses of LIP neurons. We conclude that the process of evidence accumulation leading to choice is revealed primarily in motor preparatory responses. The sensory responses exhibit a weak relationship with the animal’s behavior, but our results and simulations suggest that this relationship is likely inherited from the motor preparatory responses. We first discuss our results in the context of previous studies of area LIP and then consider their implication on the broader question of routing of information in the cortex.

### Properties of neural responses in area LIP

There has been a long debate about the relative importance of sensory salience-related signals and saccade preparatory signals in area LIP (Andersen & Buneo 2002, Barash et al 1991a, Bushnell et al 1981, Colby & Goldberg 1999). Many neurons show inherent selectivity for visual features such as direction and shape, even in monkeys that have never been trained to use such information (Fanini & Assad 2009, Sereno & Maunsell 1998). In addition, training induces stimulus selectivity that can be distinct from intrinsic selectivity (Sarma et al 2015, Toth & Assad 2002). LIP neurons also carry a rich representation of saccade plans. They display spatially selective persistent activity when the animal plans a saccade to a previously instructed, but no longer visible target (Barash et al 1991a, Gnadt & Andersen 1988). This persistent activity is dissociable from the sensory response evoked by the target (Mazzoni et al 1996) and can encode other factors that bear on the saccade plan, such as the probability that a saccade will be instructed (Janssen & Shadlen 2005) and the expected reward (Platt & Glimcher 1999, Sugrue et al 2004). The richness of saccadic planning is particularly evident in perceptual decision-making tasks, where the neuronal activity continually tracks the current state of the evidence for choosing the target in the neuron’s RF (Bollimunta et al 2012, Mazurek et al 2003).

By recording from the same LIP neurons when they belonged to the population representing either the RDM or a choice target, we could directly compare the sensory- and saccade-related responses. While both populations modulated their activity in accordance with the strength and direction of the RDM, there were important differences. This modulation was more intense when a choice target was in the RF. While the RDM elicited a strong response when it was in the RF, the dependence on direction and stimulus strength was weaker. This is unlikely to be explained by saturation of the response, because the same neurons attained higher firing rates before saccade onset when the target was in the RF (cf. Figure 3B and Figure 3C). Further, the variance and autocorrelation patterns of the neuronal responses were consistent with the predictions of noisy evidence accumulation only when the neurons contained a target in their RF. Finally, a neural correlate of decision termination was only apparent when a target was in the RF.

Although we have used the term “sensory” to describe the direction selective responses of neurons with the RDM in their RF, the gradual build-up of the firing rates of these neurons (Figure 3C) differed from the constant firing rates reported in naïve monkeys (Fanini & Assad 2009). We suspect that the responses are not sensory in the way one would characterize the responses of neurons in visual areas MT/MST or even the visual responses of LIP neurons to transient stimuli (e.g., targets) as they were remarkably slow, emerging 190 ms after stimulus onset (at the highest coherence). This is far later than the ∼50 ms latency of direction selectivity (Fanini & Assad 2009) and the ∼100ms latency for direction-category selectivity (Swaminathan & Freedman 2012), and it is longer than the 180 ms latency of decision-related signals observed in the neuronal pool representing the targets.

Together, these considerations suggest that the neuronal pool representing the RDM inherits its direction and choice related signals from the neuronal pools representing the targets. We demonstrated that a model of lateral interactions serving the general purpose of gain control (Carandini & Heeger 2011) is sufficient to produce these effects. Such lateral interactions are well established in upstream visual areas (Hunter & Born 2011, Schein & Desimone 1990, Shushruth et al 2009). In LIP, lateral interactions are thought to mediate the suppressive effect of visual stimuli presented outside a neuron’s RF (Balan et al 2008, Churchland et al 2008, Zhang et al 2017), even from distances >50° away from the RF (Falkner et al 2010, Louie et al 2011). A limitation of the present study is that we do not have access to two classes of neurons on the same trials. Recording simultaneously from neurons that represent the RDM and at least one choice target, would allow for a direct test of the lateral interactions that we modeled. For example, we would predict that the weaker leverage of the RDM-in-RF neurons would be explained away (i.e., mediated) by inclusion of Target-in-RF responses in the same GLM.

### Routing of information in cortex

We do not know how the momentary evidence represented by populations of direction selective neurons in the visual cortex makes its way specifically to the target-representing neurons in LIP. There are projections from areas MT and MST to area LIP, but it is difficult to reconcile this direct pathway with the long latency of the decision related activity in LIP. The delay of the decision related responses relative to the latency of the visual responses in LIP (∼50 ms), suggests a role for some form of memory buffer and/or a multisynaptic chain through which decision relevant information must pass before reaching the saccade planning neurons in LIP. This is one reason to suspect that apparently simple perceptual decisions may share similarities with more complex decisions that derive evidence from memory and other evaluations (Shadlen & Shohamy 2016).

We must emphasize that area LIP is not the only region that receives decision-pertinent signals in this task. Other areas involved in the planning of eye movements, such as FEF/Area 46, caudate nucleus and superior colliculus, also have access to such input (Ding & Gold 2010, Ding & Gold 2012, Horwitz & Newsome 1999, Kim & Shadlen 1999, Mante et al 2013). However, the decision related activity in these areas arises with comparable latencies, so they do not furnish an explanation for the long latency in LIP. We favor the idea that the latency is necessitated by limitations in connectivity between the many possible sources of evidence bearing on the salience of an item and the neurons that represent such items as potential affordances to the motor system. This connectivity constraint might necessitate active routing (Kastner & Pinsk 2004, Olshausen et al 1993), although this process is poorly understood.

Our results also invite caution when interpreting trial-to-trial correlations between neural response and choice behavior. The neuronal pool in LIP representing the RDM has a mean CP of 0.59, larger than the reported CP of 0.54 for neurons in area MT (Cohen & Newsome 2009) that are known to play a causal role in affecting choice and RT in this task (Ditterich et al 2003, Salzman et al 1990). One might therefore be tempted to conclude that the RDM-in-RF population plays a role in evidence accumulation leading to the decision, but this is at odds with our findings. In the RDM task, the sequential sampling framework (e.g., drift-diffusion) provides a detailed mechanistic account of evidence accumulation both at the level of behavior and at the level of its instantiation in the neural responses. This enabled us to show that only the neuronal population involved in planning of the motor action reflected the computations relevant to decision-making.

If the neurons with the RDM in the RF do not represent the evolving evidence, a natural question is what do these neurons signify? One obvious possibility is that they simply represent an object that might attract the gaze, as transient lights are wont to do. Another possibility is that they represent the focus of spatial attention (Colby & Goldberg 1999). However, this focus should be initially on the RDM and then either remain stationary through the decision or gradually give way to the chosen target. This is inconsistent with the dynamics observed in our data, which look like a muted version of the decision related signals exhibited by neurons with a choice target in the RF. The same objection applies to the proposal that these neurons represent the salience of the RDM (Bisley & Goldberg 2010). A more speculative idea is that the neurons that contain the RDM in their RF confer information bearing on the spatial origins of the evidence—that is, they help to bind the location of the thing we are deciding about to the decision itself, which is about what to do.

## Acknowledgements

This work was supported by HHMI, NEI (R01 EY11378), NCRR (RR00166). The authors thank Drs. Fetsch, Jeurissen, So, Steinemann and Zylberberg for advice on earlier versions of the manuscript.

## FIGURE LEGENDS

**FIGURE 2-EXTENDED DATA 1:**
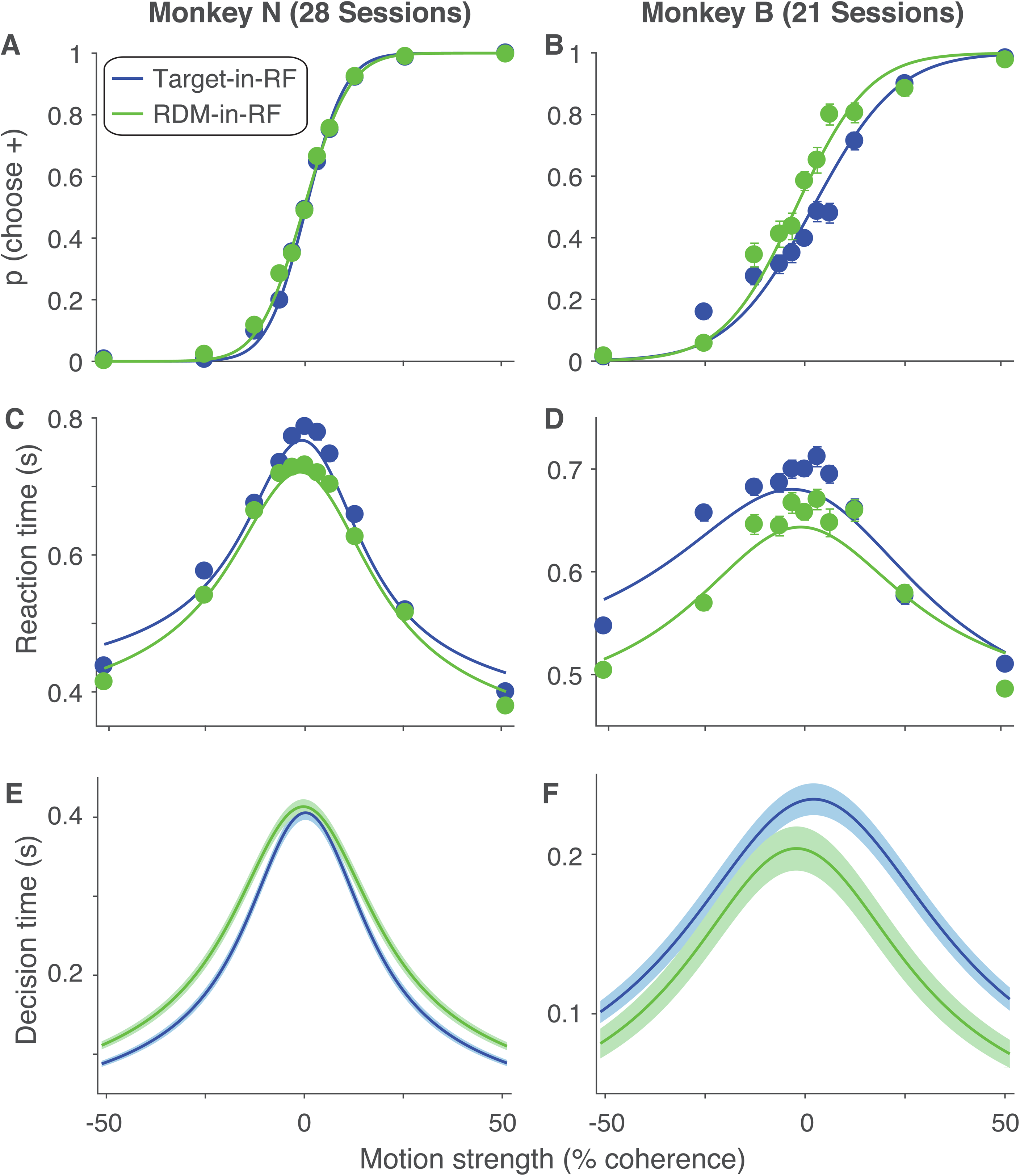
Simultaneous fit of RT and choice with a diffusion-to-bound model. The probability of choosing the positive direction target **(*A,B*)** and the mean RTs **(*C,D*)** are plotted as a function of motion strength and direction (indicated by sign of coherence; see Methods) for the two monkeys in the two stimulus configurations. The curves are fits to the data from a diffusion-to-bound model with nonstationary bounds (see Methods). The fit parameters are shown in Table 1. ***E,F:*** Mean decision times (solid curves) derived from the model fits, plotted as a function of motion strength. Shading is ±1 S.E.

**FIGURE 3-EXTENDED DATA 1 and 2:**
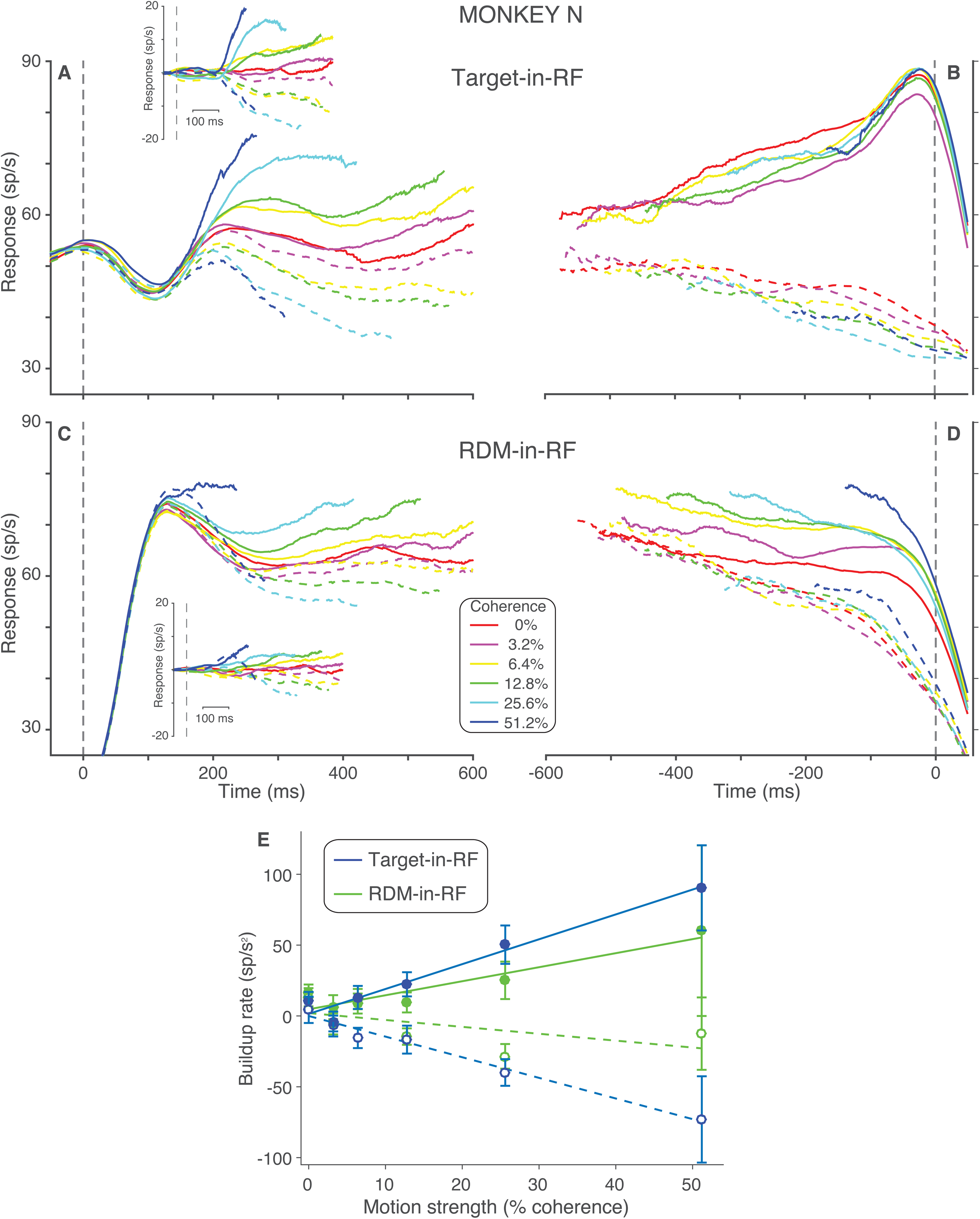
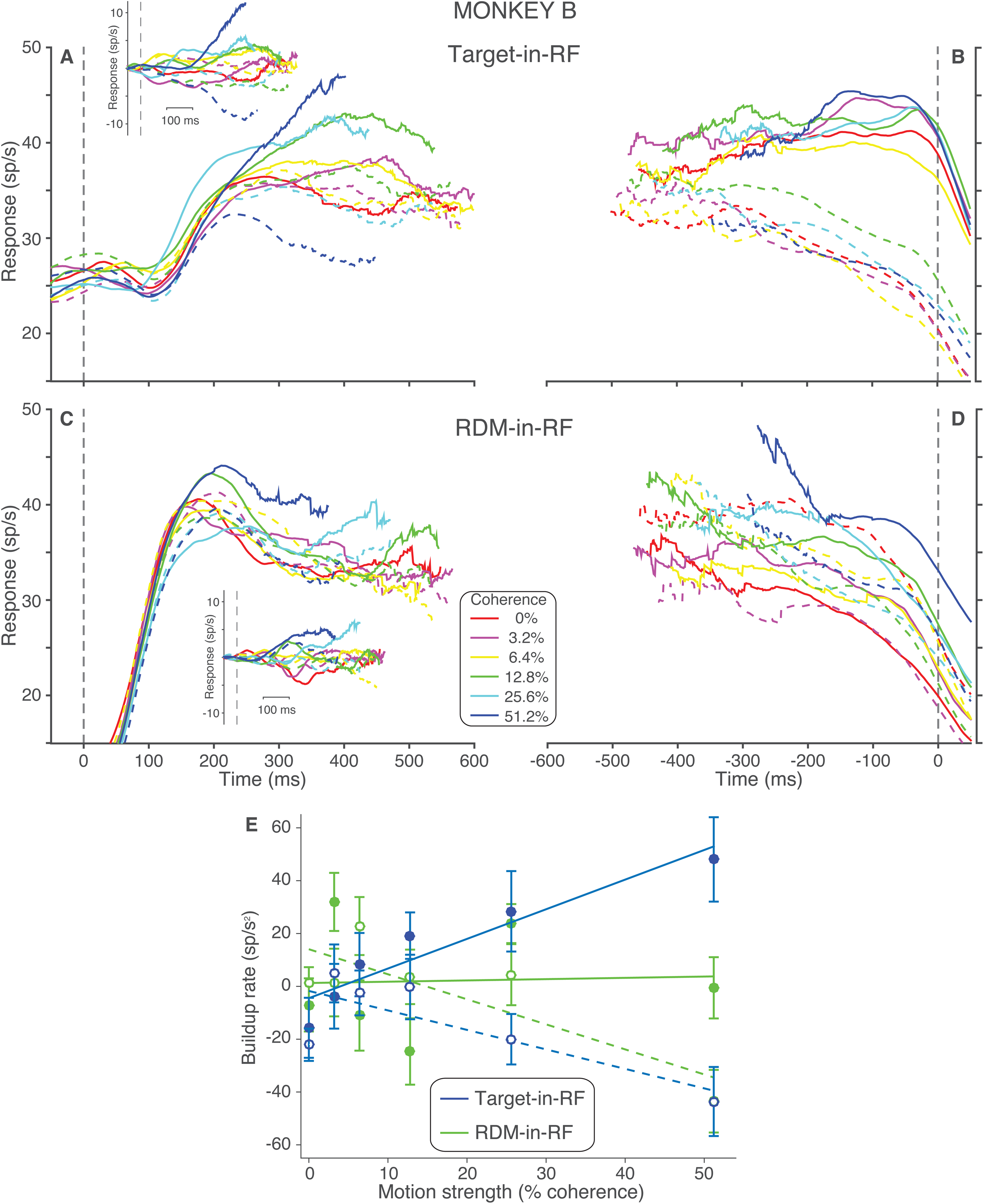
Population responses of neurons in individual animals. Neural responses that were shown in Figure 3, panels A-D and F plotted from data pooled separately for each individual monkey.

